# Sequence determinants of human gene regulatory elements

**DOI:** 10.1101/2021.03.18.435942

**Authors:** Biswajyoti Sahu, Tuomo Hartonen, Päivi Pihlajamaa, Bei Wei, Kashyap Dave, Fangjie Zhu, Eevi Kaasinen, Katja Lidschreiber, Michael Lidschreiber, Carsten O. Daub, Patrick Cramer, Teemu Kivioja, Jussi Taipale

## Abstract

DNA determines where and when genes are expressed, but the full set of sequence determinants that control gene expression is not known. To obtain a global and unbiased view of the relative importance of different sequence determinants in gene expression, we measured transcriptional activity of DNA sequences that are in aggregate ∼100 times longer than the human genome in three different cell types. We show that enhancers can be classified to three main types: classical enhancers^1^, closed chromatin enhancers and chromatin-dependent enhancers, which act via different mechanisms and differ in motif content. Transcription factors (TFs) act generally in an additive manner with weak grammar, with classical enhancers increasing expression from promoters by a mechanism that does not involve specific TF-TF interactions. Few TFs are strongly active in a cell, with most activities similar between cell types. Chromatin-dependent enhancers are enriched in forkhead motifs, whereas classical enhancers contain motifs for TFs with strong transactivator domains such as ETS and bZIP; these motifs are also found at transcription start site (TSS)-proximal positions. However, some TFs, such as NRF1 only activate transcription when placed close to the TSS, and others such as YY1 display positional preference with respect to the TSS. TFs can thus be classified into four non-exclusive subtypes based on their transcriptional activity: chromatin opening, enhancing, promoting and TSS determining factors – consistent with the view that the binding motif is the only atomic unit of gene expression.

## Introduction

The temporal and spatial pattern of gene expression is encoded in the DNA sequence; this information is read and interpreted by transcription factors (TF), which recognize and bind specific short DNA sequence motifs^2, 3^. Major efforts have been undertaken to determine the DNA binding specificities of TFs *in vitro*^4,5,6,7,8^ and in mapping their binding positions *in vivo*^9,10,11^. TFs regulate gene expression by binding to distal enhancer elements, and to promoters located close to the transcription start site (TSS)^1, 10, 12^. Both enhancers and promoters are characterized by RNA transcription^13, 14^, presence of open chromatin^15, 16^ and histone H3 lysine 27 acetylation^17^ (H3K27ac). In addition, promoters and enhancers are marked by histone H3 lysine 4 (H3K4) trimethylation and monomethylation^18^, respectively. Although these features can be mapped genome-wide in a high-throughput manner, they are correlative in nature and do not establish that an element can act as an enhancer. To more directly measure enhancer activity, several investigators have developed massively parallel reporter assays (MPRAs) and used them to study the activity of yeast^19, 20^, *Drosophila*^21, 22^ and human^23,24,25,26,27,28^ gene-regulatory elements on a genome-wide scale. However, unbiased discovery of sequence-determinants of human gene expression by only analyzing genomic sequences is made difficult by the fact that the genome is repetitive and evolved to perform multiple functions in addition to transcription. Furthermore, the human genome is too short to even encode all combinations, orientations and spacings of the 1639 TFs^2^ in multiple independent sequence contexts. Thus, despite the vast amount of information generated by the genome-scale experiments, most sequence determinants that drive the activity of human enhancers and promoters, and the interactions between them remain unknown.

## Results

### Ultra-complex MPRA libraries with 100 times human genome coverage

To comprehensively characterize the sequence determinants of human gene regulatory element activity, we developed a set of four MPRA libraries that cover more than 100 times the sequence space of the human genome (see **Methods** for details). The libraries are based on the STARR-seq design^21^, in which putative enhancers are cloned to an exon, and the enhancer activity is then read using RNA-sequencing (**Fig. 1a**). Three libraries were designed to measure enhancer activities of (i) combinations of known TF binding motifs, (ii) ∼500 bp fragments of genomic DNA and (iii) synthetic random 170 bp sequences, and the fourth library was designed (iv) to measure both enhancer and promoter activities of synthetic random 150 bp sequences. To enable analysis of the effect of DNA methylation on transcriptional activity, we developed a MPRA vector that is devoid of CG dinucleotides: in designs i to iii, the Lucia reporter gene is driven by CG depleted minimal promoters, whereas in design iv, the promoter is replaced by 150 bp random DNA sequences (**Fig. 1a**; see **Methods** and **Tables S1, S2**). Sequencing of the input libraries revealed their ultra-high complexity reaching billions of unique fragments (**Fig. S1a,b**; see **Methods).**

**Fig. 1.**
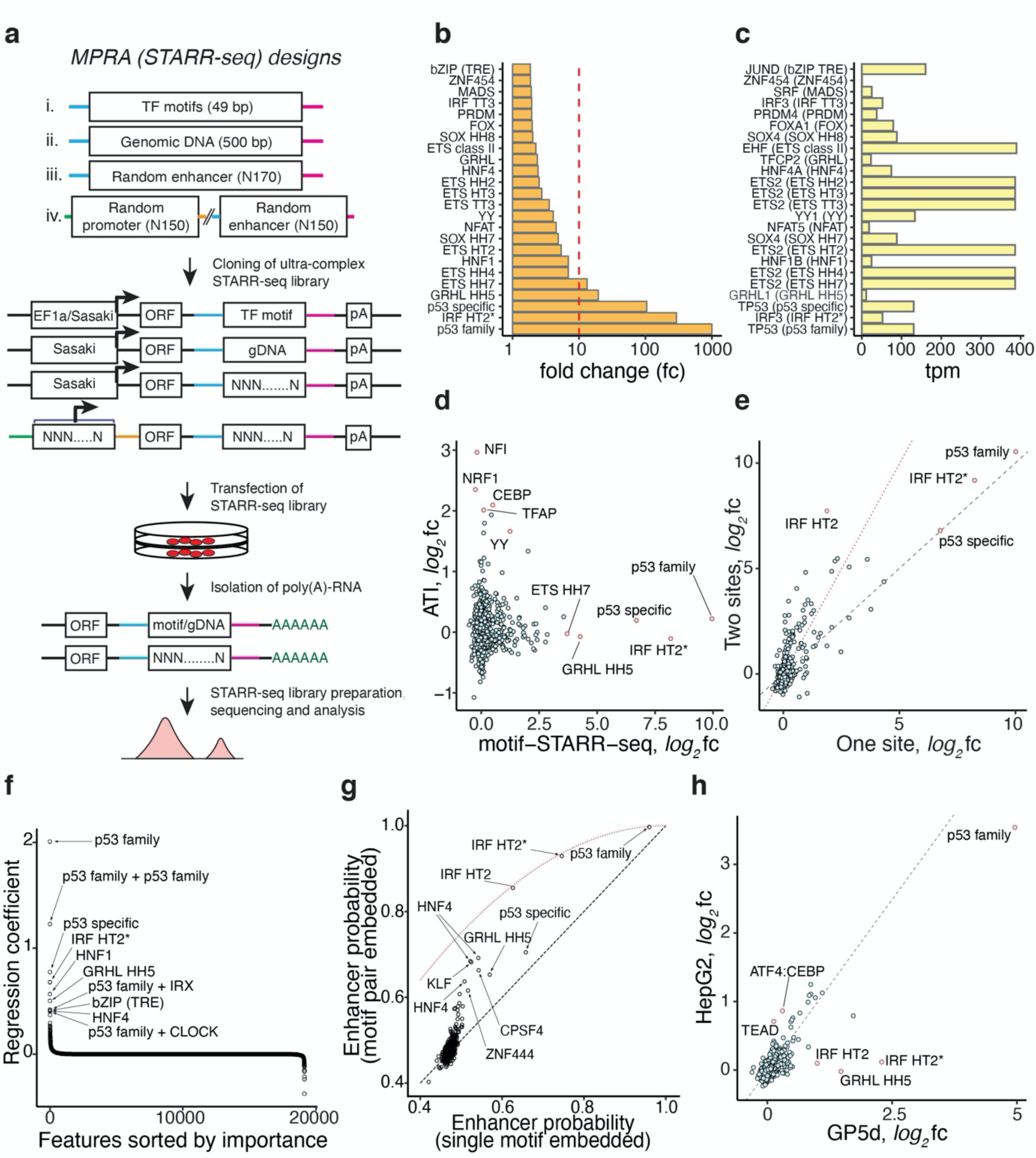
Few TFs display strong transcriptional activity in cells. **a,** Schematic representation of the MPRA (STARR-seq) reporter construct and its variations used in the study. Design of the MPRA reporter constructs and experimental workflow for measuring promoter or enhancer activity in mammalian cells is shown. For enhancer activity assays, DNA library comprising of either synthetic TF motifs (i), human genomic fragments (ii), or completely random synthetic DNA oligonucleotides (iii) is cloned within the 3’UTR of the reporter gene (ORF) driven by a minimal δ1-crystallin gene (Sasaki) promoter or CpG-free EF1α promoter. For random promoter and random enhancer (iv) activity assays random synthetic DNA sequences are cloned upstream of the ORF in place of the minimal promoter and downstream of the ORF in the 3’ UTR. The motif library (i) comprises a total of 92,918 individual sequence patterns including 1121 TF consensus sequences^6, 8^ and 30,123 mutant consensus sequences in two motif-depleted sequence contexts (to control for effect of flanking bases). Each pattern only contained one type of consensus sequence, present once, twice and/or three times. Multiple consensus sequences were arranged in different spacings and orientations relative to each other (see **Methods** and **Tables S3, S4** for details). In the experiments, the MPRA reporter libraries are transfected into human cell lines and total RNA is isolated after 24 hours of transfection followed by enrichment of reporter-specific RNA and Illumina library preparation, sequencing and data analysis. The active promoters are recovered by mapping their transcribed enhancers to the input DNA and taking the corresponding promoter pairs. **b,** Enhancer activity of HT-SELEX motifs. Synthetic TF motif library was transfected into GP5d cells and motifs were analyzed for enhancer activity. Median fold change (fc) of the sequence patterns containing once the motif consensus or its reverse complement over the input library is shown. Red dotted line marks 1% activity related to the strongest motif. Dimeric motifs are indicated by orientation with respect to core consensus sequence (GGAA for ETS, ACAA for SOX, AACCGG for GRHL and GAAA for IRF; HH head to head, HT head to tail, TT tail to tail, followed by gap length between the core sequences). Asterisk indicates an A rich sequence 5’ of the IRF HT2 dimer. **Table S5** describes the naming of the motifs in each figure. **c,** Expression level (tpm; reads per transcript per million) in GP5d cells for TFs that bind to the motifs with strong activity in panel (**b)**. The motif that the TF can bind is shown in parentheses. Note that the order is the same as in panel (**b)**. **d,** Comparison between motif transcriptional activity (x-axis) and biochemical binding activity from ATI assay (y-axis) in GP5d cells. Note that the Pearson correlation between the transcriptional activity and the biochemical activity is relatively low (R=0.032) indicating that the TF motifs responsible for the activities are largely distinct. Note also that some TFs can bind to two or more slightly different motifs that can have somewhat different activities (e.g. p53-family motif is bound by all p53 family members, p53, p63 and p73, whereas the p53-specific motif is only bound by p53). **e,** Analysis of the effect of number of TF binding sites on enhancer activity from synthetic motif library in GP5d cells. For each TF motif, the fold change (log_2_) compared to the input is shown for one versus two binding sites. The dashed line represents the expected fold change if two sites have the same effect as one, and the red dotted line represents the expected fold change if the two sites act in an additive manner. Note that additive effect in the logarithmic space equals multiplication of fold changes. **f,** Regression coefficients for different TFs and TF pairs from logistic regression analysis of enhancer activities measured from random enhancer library in GP5d cells (see **Methods** for details). Coefficient value (y-axis) is shown for all TFs and pairs of TFs, with features with the strongest predictive power indicated by labels. **g,** Non-linear effect of multiple motifs inserted into sequences scored using the CNN trained on GP5d random enhancer STARR-seq data. Note that a pair of the same motifs (indicated by labels) increases the predicted enhancer probability of the sequence above that expected from a single motif (dashed black line), but not above that expected from a model assuming independent binding to two motifs in the same sequence (red dotted line). Note that CNN identifies similar features than the enrichment analysis shown in **Fig. S4a**. **h,** Comparison of enhancer activity of motifs measured from random enhancer library in two mammalian cell lines, GP5d colon cancer and the HepG2 liver cancer cell lines, showing the fold change of motif match count over input in each cell line (black dashed line indicates identical activity between the cell lines).

### Few TFs display strong transcriptional activity in cells

To measure the enhancer activity of the known TF consensus sequences, we transfected GP5d colon carcinoma cells with the motif libraries (**Fig. 1a**, i), and purified total poly(A+) RNA from the transfected cells. The synthetic motif sequences that were transcribed to RNA were recovered using RT-PCR and the abundance of each sequence then quantified by massively parallel sequencing (see **Methods**). Comparison of the median activities of the individual TF consensus sequences revealed that several TFs had enhancer activity in GP5d cells (**Fig. 1b**; **Fig. S2a**; **Table S5**). The most active motifs displayed similar activities when placed in different sequence contexts, and between experiments using two different basal promoters, δ1-crystallin and CpG-free EF1α promoters (**Fig. S2a,b**). Although the method used measures activities of consensus sequences and/or motifs, the TFs or groups of TFs that bind to the sequences can be inferred from the motifs and TF expression levels^29^. Such combined analysis revealed that TFs that express at a relatively high level in GP5d cells can bind to the strongly active motifs (average 102 transcripts per million (tpm) vs. 26 tpm for all genes and 20 tpm for all TFs from ref.^2^); however, the correlation between motif activity and expression of corresponding TFs was weak (**Fig. 1c**), indicating that expression alone does not determine transcriptional activity. The consensus sequence corresponding to the p53 protein family (p53, p63 and p73) displayed the strongest enhancer activity in this assay (**Fig. 1b**). As the library contained each single-base substitution to the p53 family consensus sequence, we were able to generate an activity position weight matrix (PWM) for the consensus. The activity PWM was highly similar to the SELEX-derived motif for the p53 family (**Fig. S2c**), indicating that the measured enhancer-activity originated from a p53 family TF, and that the assay can be used to faithfully measure TF activities in cells.

Quantitative analysis revealed that only three other motifs, representing interferon regulatory factor (IRF), grainy-head like (GRHL) and E26 transformation-specific (ETS) TFs had activity that was within 1% of the maximal activity observed for the p53 family motif (**Fig. 1b**, dotted line). Of note, the two most enriched motifs, p53 and IRF, are bound by TFs that respond to cellular alarm signals such as DNA damage (p53) and cytoplasmic DNA (IRF), suggesting that ordinary transfection can induce these alarm signals.

Comparison of enhancer activities of motifs with the DNA binding activities of respective TFs measured by active TF identification (ATI) assay^29^ revealed that the transcriptional and DNA binding activities were only weakly correlated (log_2_ fold change, Pearson R^2^= 0.032), with only a subset of motifs with moderate enhancer activity (e.g. YY) also displaying moderate DNA binding activity (**Fig. 1d**).

### Synergy, additivity and saturation of activity

Apart from simple cellular alarm signals, most transcription is thought to require combinatorial action of many TFs^30,31,32^. Consistently with this, we observed that the average activity of all consensus sequences was very low, and for the majority of the TFs, the enhancer activity increased as a function of the number of consensus sequences (**Fig. S2d**, red horizontal lines). Conversely, for the TFs that can activate transcription alone (e.g. p53, IRF HT2 with A-rich 5’ sequence), two consensus sequences had lower activity than that predicted from an additive model (**Fig. 1e**, red dotted line), presumably due to saturation of both the occupancy and the downstream transcriptional activation. For TFs with intermediate activity levels (e.g. NFAT and YY), activity increased linearly rather than synergistically as a function of the number of binding sites (**Fig. S2d**). The simplest model consistent with these observations is that human enhancer activation requires overcoming a repressive activity, after which activation is linear (additive) until it starts to saturate as it approaches a maximum level.

### Transcriptional enhancers can readily evolve *de novo* from random sequences

To discover sequence features that contribute to human enhancer activity in an unbiased manner, we measured the activity of sequences from the extremely complex random enhancer library (**Fig. 1a**, iii) in GP5d cells. Motif-mapping across replicate experiments indicated that motif activities were highly reproducible (**Fig. S3a**); we also observed a linear increase in enhancer activity as a function of the number of motif matches for TFs with moderate activity (e.g. bZIP TRE), and a saturation of the enhancing effect of multiple matches of p53 family motifs (**Fig. S4a,b**). Importantly, we also detected enrichment of motifs corresponding to known TFs specific to colon cancer and intestinal lineage, such as TCF/LEF, GRHL, and HNF4 (**Fig. S4a**). *De novo* motif mining identified several TF motifs; most of these were for individual TFs or conventional heterodimers, suggesting that the backbone of the transcriptional system is based on individual TFs acting together without strict spacing preferences or grammar. However, one strong *de novo* motif identified was for a new ETS-bZIP composite motif, revealing a potential role for ETS-bZIP combinatorial control in colon cancer cells (**Fig. S5a**). These results indicate that transcriptional enhancers are relatively simple low information content sequence features that can be evolved from random sequence in a single enrichment step.

To determine all sequence features present in the *de novo* evolved enhancers, we used machine learning classifiers, with 70%, 15% and 15% of the data used for training, validation and test sets, respectively. First, we determined the importance of known motif features using a logistic regression model (see **Methods**); we found that only a handful of known TF binding motifs are needed for optimal logistic regression classification (**Fig. 1f**; **Fig. S4c**; see **Methods** for details) of the active enhancer sequences from the inactive ones, and that their interactions were largely additive, as specific pairwise combinations did not add substantially to the predictive power. The most predictive features were motifs for known TFs important for tumorigenesis and for colon development (**Fig. 1f**). Next, to identify all sequence features, we trained a convolutional neural network (CNN)-based classifier similar to DeepBind^33^ on the sequence data. This method is capable of learning the sequence motifs, their combinations, and their relative weights *de novo*. The CNN classifier performed substantially better than logistic regression using the same training, validation and test sets (**Fig. S4c,d**; see **Methods** for details). Analysis of the CNN classifier revealed that it had learned motif features similar to those identified by the logistic regression (**Fig. 1g**). No other motifs were detected, suggesting that the CNN can either improve the PWM motifs themselves, or that there are other types of sequence features, or interactions between features (**Fig. 1g**) that are important in classification that are not captured by the logistic regression model (see **Methods**).

### Transcriptional landscape of a cell is dominated by housekeeping TFs

To determine whether enhancers are similar between cell types, we used the random enhancer library (**Fig. 1a**, iii) to identify sequence features important for enhancer activity in HepG2 hepatocellular carcinoma cells. Comparison of enhancer motifs between the GP5d and HepG2 cells revealed that most motifs had similar enhancer activity across the cell lines (**Fig. 1h**). The motifs that had differential activity corresponded to lineage-determining TFs (GRHL in GP5d), TFs important for tissue function (TEAD and ATF4:CEBPB in HepG2), or were related to the known deficiency of the HepG2 cells in interferon signaling (IRF3)^34^. Importantly, the lineage-determining factors showed clear differential expression between the two cell types (**Fig. S6**), indicating that activities of individual TFs are commonly affected by expression level, despite the differences in specific activities of TFs leading to a low correlation between expression and activity across all TFs (see **Fig. 1c**). Taken together, the transcriptional landscape of a cell is dominated by cell-biological or “housekeeping” TFs; the strongest differences between cells represent known TFs that are important for specification and/or function of the specific lineages.

### Comparison of genomic enhancers and *de novo* evolved enhancers

To determine how sequence features combine to generate functional genomic enhancers, we assayed enhancer activity in GP5d and HepG2 cells using ∼500 bp fragments derived from GP5d genome (**Fig. 1a**, ii) before and after methylation of the library. To determine the role of p53 in enhancer activity, we performed similar experiments also in p53^-/-^ GP5d cells (**Fig. S7a-c**). The high complexity of the library (2.09 × 10^9^ distinct clones; **Fig. S1**), replicate concordance and excellent signal-to-noise ratio allowed detection of enhancer activity at ∼1.5 bp resolution (**Fig. S3c,d**; see **Methods**). Consistent with the known association between accessible chromatin, TF binding and enhancer activity, the STARR-seq peaks overlapped significantly with chromatin accessibility (**Fig. 2a-c**; **Fig. S7d-g**). Furthermore, ATAC-seq peaks could be predicted by a CNN trained using genomic or random STARR-seq sequences (**Fig. S7i**), indicating that the sequence features discovered using STARR-seq correspond partially to the features that are associated with open chromatin *in vivo*. However, the overlap between ATAC-seq and STARR-seq was only partial; detailed analysis (**Fig. 2a,c**) revealed six classes of elements: i) closed chromatin enhancers (STARR-seq+, ATAC-seq-), ii) cryptic enhancers (silenced STARR-seq+ regions), iii) promoters (ATAC-seq+ with or without STARR-seq), iv) chromatin-dependent enhancers (STARR-seq-/low, ATAC-seq+ with active histone marks), v) structural chromatin elements (STARR-seq-, ATAC-seq+, CTCF+) and vi) classical enhancers (STARR-seq+, ATAC-seq+).

**Fig. 2.**
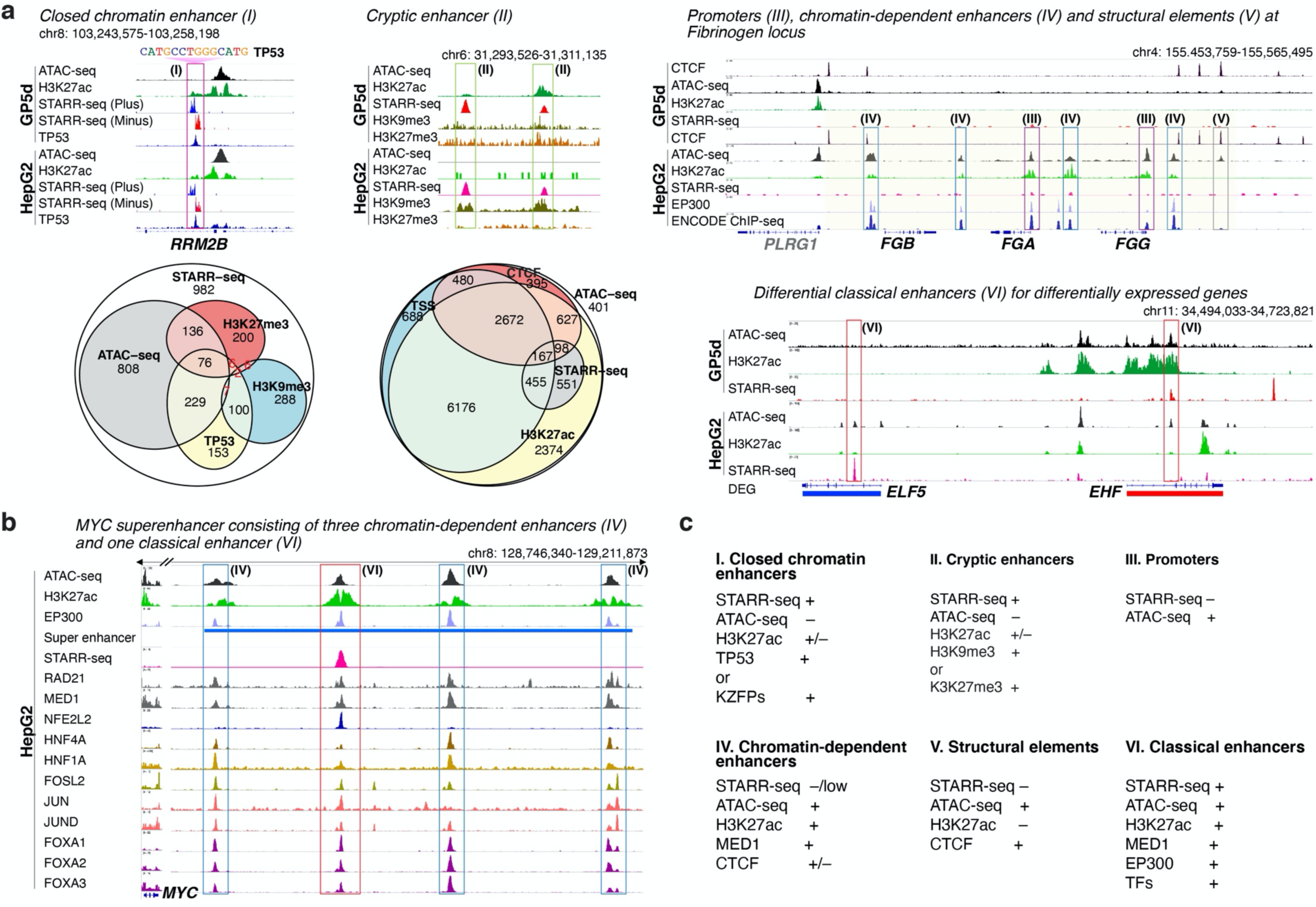
Genomic STARR-seq reveals six types of regulatory elements. **a**, Six types of regulatory elements classified on the basis of STARR-seq signal and chromatin features such as accessibility (ATAC-seq), TF binding, and epigenetic modifications. Euler diagrams (bottom) show the overlap between genomic STARR-seq peaks and different genomic features (left) and between ATAC-seq peaks and different genomic features (middle) in HepG2 cells. Some of the small intersections are not shown, see **Fig. S7g** for the full list. Genome browser snapshots showing examples of different types of regulatory features in HepG2 and GP5d cells are also shown. Colored boxes marked with roman numerals correspond to the different types of elements listed in panel **c**; clockwise from top: closed chromatin enhancer (**I**) devoid of H3K27ac or ATAC-seq signal at TP53-target gene *RRM2B*, both plus and minus strand STARR-seq signal is shown; cryptic enhancer (**II**) overlapping with repressive histone marks; promoters (**III**) and chromatin-dependent enhancers (**IV**) and structural CTCF element (**V**) at the fibrinogen locus, the Encode ChIP-seq track includes combined ChIP-seq signal for 206 TFs^11^; tissue-specific classical enhancers (**VI**) detected for *ELF5* (higher expression in HepG2, blue) and for *EHF* (higher expression in GP5d, red). Note that the STARR-seq peaks are specific to the cell types where the adjacent gene is expressed. **b**, Chromatin-dependent enhancers and classical enhancers combine to form super-enhancers. Genome browser snapshot of a MYC super-enhancer in HepG2 cells marked by STARR-seq peak overlapping with the binding site for TF with strong transactivation activity (NFE2L2) converging on equidistant chromatin enhancers bound by cohesin, mediator, forkhead and other liver specific TFs. **c**, Summary of the features that define the six genomic element types.

All three types of enhancer (closed chromatin, chromatin dependent and classical) appeared active based on the fact that inclusion of the corresponding features improved prediction of differential gene expression between GP5d and HepG2 cells (**Table S6**; see **Methods**). Analysis of ChIP-seq peaks and motifs present in the different classes of elements revealed that classical and closed-chromatin enhancers bound to TFs and contained motifs that were similar to those that were found in active elements selected from random sequences (**Fig. S5b**; see also **Fig. 1h**). Chromatin-dependent enhancers contained an additional set of motifs that were not present on random STARR-seq, including motifs for FOXA and HNF4A. These results indicate that cells contain three distinct classes of enhancers. One of these is the classical enhancer described by Banerji et al. (ref.^1^); these elements contain open chromatin and transactivate a heterologous promoter regardless of position or orientation. The other main type of cellular enhancer appears dependent on chromatin and cannot be effectively detected using STARR-seq (see also ref.^35^); these elements are associated with strong signal for open chromatin and the activating histone mark H3K27ac. Conversely, closed chromatin contains a third type of enhancer activity that cannot be detected using these classical marks, and whose detection requires STARR-seq.

Consistently with few TFs determining the overall transcriptional landscape of a cell, the genomic STARR-seq peaks were enriched for relatively few motifs. The motifs themselves were similar to known monomeric, dimeric and composite TF motifs determined using HT-SELEX^6^ and CAP-SELEX^36^ (**Fig. S5a**). The motifs discovered from genomic and random enhancers were largely similar (**Fig. S5a**). The main difference was the discovery of the pioneer factor FOXA from genomic fragments, suggesting that although FOXA proteins do not strongly activate transcription, their motifs are associated with classical genomic enhancers because of the ability of FOXA to displace nucleosomes and/or to open higher order chromatin. This is also consistent with the fact that FOXA motifs were strongly enriched in chromatin-dependent enhancers that lacked STARR-seq signal (**Fig. S5b**). Many of the motifs discovered from genomic STARR-seq also displayed strong DNA binding activity in an ATI assay (**Fig. S5a**), indicating that strong DNA binders are important for *in vivo* enhancer activity, potentially because they are capable of opening chromatin^29^. In summary, the sequence features of classical genomic enhancers are highly similar to those evolved from random sequence; these motifs define the classical enhancer activity of a cell. In addition to this activity, additional chromatin-dependent enhancers confer tissue-specificity to genes; these elements are characterized by motifs for TFs that have lower transactivation activity, suggesting that these TFs act via chromatin to facilitate the activity of promoters and associated classical enhancers. Consistently with this view, the strongest cellular enhancers, super-enhancers, typically consist of arrays of chromatin-dependent elements associated with a classical enhancer (**Fig. 2b**; **Fig. S7h**).

### Sequence features of promoters and enhancers evolved from random sequences

To identify sequence determinants of human promoter activity, we assayed the activity of the “binary STARR-seq” library consisting of random sequences placed in the position of both the promoter and the enhancer (**Fig. 1a**, iv). For this analysis, we used two tumor cell lines (GP5d and HepG2) and an untransformed cell line derived from retinal pigment epithelium (RPE1). Robust promoter activity was observed in all three cell lines from a subset of the random sequences, and motif-mapping across replicate experiments in GP5d cells showed that motif activities were highly reproducible (**Fig. S3b**). As observed for the motifs at active enhancers, most motifs enriched at promoters were similar in all cell types. The motifs that displayed differential activity were linked to lineage determination (e.g. HNF1A) and specialized cell functions (ATF4:CEBP in HepG2; **Fig. 3a**). Comparison of the evolved sequences in GP5d cells revealed that many sequence motifs were enriched in both the promoter and the enhancer positions (**Fig. 3b**). However, elements with preferential enrichment were also detected. For example, while p53 and YY motifs were similarly enriched at promoters and enhancers, ETS motifs were preferentially, and NRF1 motifs almost exclusively enriched at promoters (**Fig. 3c**). No motif enriched only at enhancers, indicating that all motifs with enhancing activity can also act from a proximal position at the promoter (**Fig. 3b**).

**Fig. 3.**
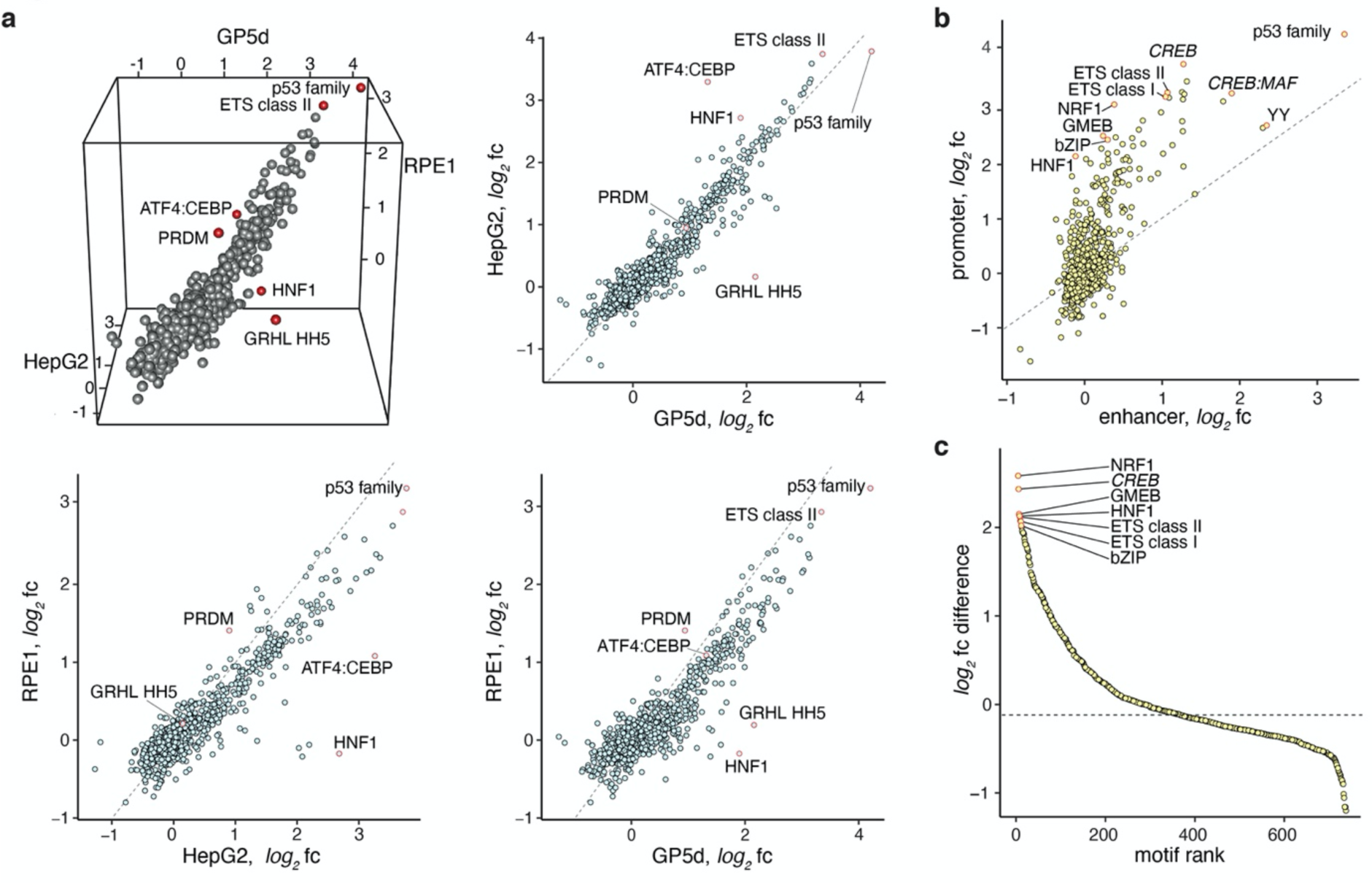
Comparison of sequence features of *de novo* evolved human promoters and enhancers. **a,** Plot showing the enrichment of TF motif matches in promoters selected from completely random sequences across three mammalian cell lines: GP5d colon cancer, HepG2 liver cancer, and RPE1 retinal pigmented epithelial cells (dashed line marks identical activity). Dimeric motifs are indicated by orientation with respect to core consensus sequence as described in legend to Fig. 1b. **b,** Comparison between enrichment of motif matches at enhancers (x-axis) vs promoters (y-axis) in GP5d cells (active sequences selected from random promoter and enhancer sequences). The motifs marked with italic typeface are *de novo* motifs mined from the GP5d TSS-aligned sequences. **c,** Motifs that enrich specifically in the promoter position, as measured by a difference in log_2_ fold change. The motifs that are enriched the most are indicated by red circles and labeled. The motifs marked with italic typeface are *de novo* motifs mined from the GP5d TSS-aligned sequences. Note also that the motifs with negative difference in log_2_ fold change (below dotted line) are repressive and decrease promoter activity; no motif specifically enriches at enhancers (see panel **b**).

### A novel G-rich element that interacts with the TSS

To evaluate the positioning of the different features relative to the TSS, we first determined the TSS position within the promoters derived from random sequences by recovering the 5’ end of the transcript using a template switch (**Fig. 4a**), yielding 85,217 unique TSS positions. The obtained reads preferentially mapped to positions that displayed a 10 bp periodicity relative to the STARR-seq vector (**Fig. S8a**). The pattern was consistent with the loading of the RNA polymerase II pre-initiation complex onto the random sequence, as the periodic pattern weakened when the ∼100 bp region occupied by the pol II complex included plasmid-derived constant sequences. Similar, but very weak periodicity was also observed in p53 motif positioning at the enhancer, suggesting that plasmid supercoiling, or some sequence feature in the vector makes one side of the DNA more accessible (**Fig. S8b**).

**Fig. 4.**
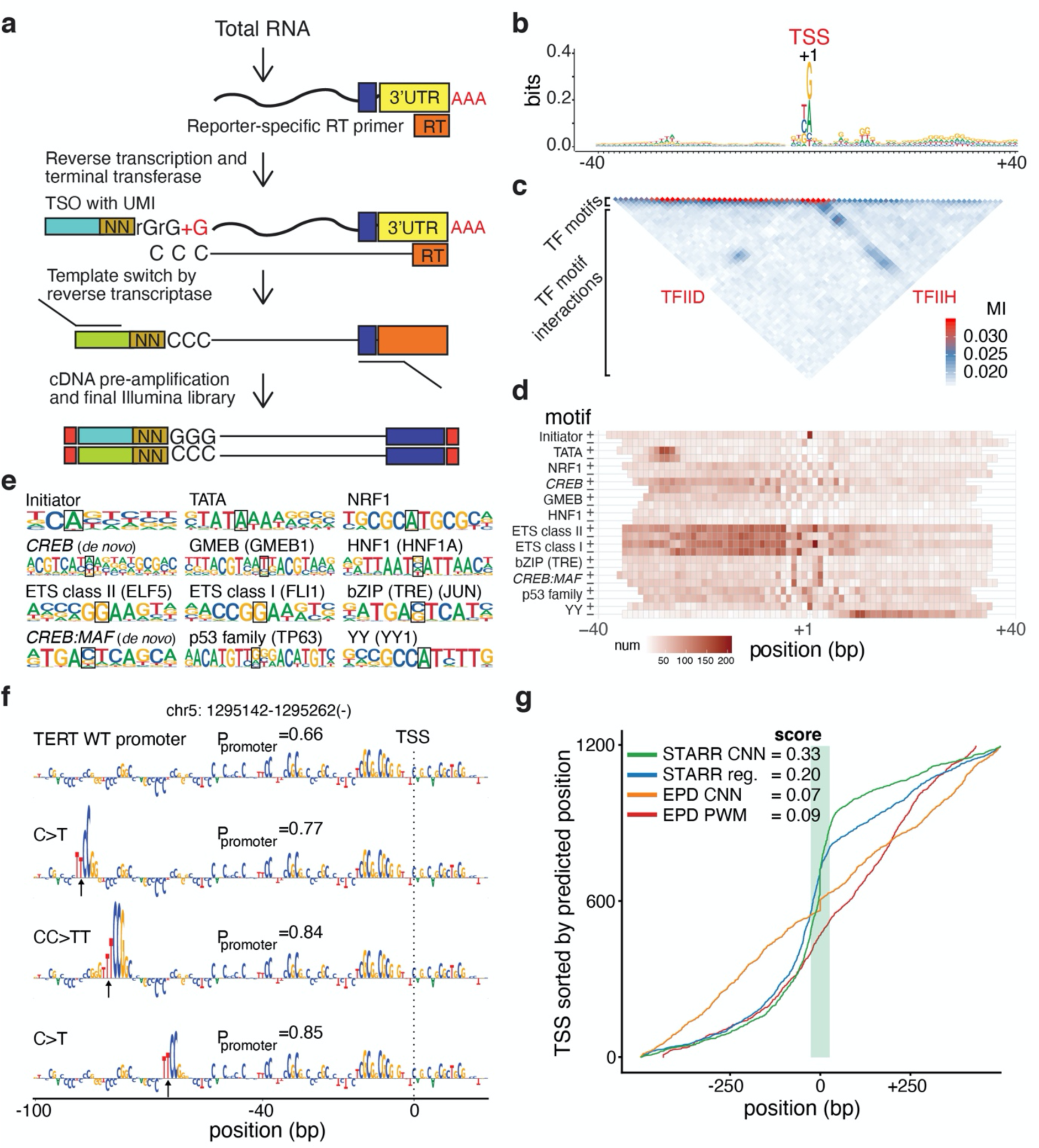
Analysis of positional specificity of sequence elements defining human promoters. **a,** Cartoon showing the design of the template-switch chemistry used to capture the 5’ sequence of the transcribed RNA using a template switch oligo (TSO) to determine the precise location of the transcription start site (TSS) within the random DNA sequences cloned at the place of the promoter. **b,** Sequence logo constructed from the 50,787 evolved GP5d promoter sequences aligned based on the measured position of their TSS (+1). **c,** Mutual information (MI) plot from the random promoter measurements. Note that most MI is observed close to the diagonal (indicating presence of TF motifs), but that two longer-range interactions are also observed, one between the TATA-box and TSS, and the other between the TSS and a G-rich element 3’ of it. **d,** Heat map showing positional preferences of the TF motifs from panel (**e**). Heat map color indicates the number of matches for the motif in one strand (p-value cut-off 5 × 10^-4^). The motifs marked with italic typeface are *de novo* motifs mined from the GP5d TSS-aligned sequences. **e,** Sequence logos of the motifs shown in the heat map (**d**). The information content center column used to position the matches in the heat map is highlighted. The motifs marked with italic typeface are *de novo* motifs mined from the GP5d TSS-aligned sequences. **f,** CNN predictor correctly identifies cancer-associated mutations in the TERT promoter. Top: predicted sequence determinants at the TERT promoter as determined by DeepLIFT^73^ analysis of the CNN. Bottom: the effect of three different driver mutations^43,44,45^ on predicted activity of the TERT promoter (the exact mutated bases highlighted with arrows). Note that the predictor identifies the ETS motifs that are generated by the driver mutations and the predicted promoter probabilities (P_promoter_) are higher for the mutant promoters. **g,** CNN trained on random promoter data outperforms PWM-based models, regression models, and CNN trained on genomic promoter data in predicting active TSS positions in GP5d cells. The cumulative distance of the predicted TSS positions from the annotated is shown against a test set of GP5d genomic TSSs (∼1200 sequences) for CNN trained on human genomic TSS data (orange), and for PWM based model (red), regression model using positional match data (blue) and CNN (green) trained on promoters evolved from random sequences. The genomic TSS positions are all aligned at 0; the curves mark predicted TSS positions for each model, sorted by distance from the annotated TSS position. The score indicates the fraction of predicted TSS positions falling within ±25 bp (the area shaded with green) from the annotated TSS positions in the genome for each model separately. Models trained on STARR-seq promoters evolved from random sequences are better at finding the correct annotated TSS position in the genome than models trained on genomic promoter sequences.

Alignment of the recovered sequences with respect to the TSS positions (see **Methods**) revealed a relatively high information content feature located at the TSS that corresponded to the classic Initiator motif (**Fig. 4b**). In addition, a clear AT-rich region was observed at the canonical -30 position of the TATA-box. However, we did not detect other TSS-proximal motifs that have previously been described (BRE, DPE, MTE, DCE, X-core promoter element, and TCT^32, 37,38,39.40^). The transcript side was characterized by a modest increase in G across a relatively wide region (+10 to +35); this, to our knowledge novel feature is also observed in genomic promoters (**Fig. S8c**). To identify interactions between the features, we performed mutual information analysis^41^. Strongest signal was for very short-range interactions located 5’ to the TSS; this signal represents enrichment of individual TF motifs. Two mutually exclusive longer-range interactions were detected, one between the TATA-box and the TSS, and the other between the TSS and the G-rich downstream sequence (**Fig. 4c**). This pattern is consistent with the loading of the RNA polymerase II either “heel first” (TFIID) or “toe first” (TFIIH) with respect to the TSS.

Motif mapping revealed that many TF motifs were also specifically positioned and oriented relative to the TSS (**Fig. 4d,e**). The strongest positional signals were observed for the TATA-box, Initiator and YY (YY1). YY1 motifs were mainly enriched on the transcript side (the first C of the CCAT sequence occurring on the - strand at position +12), oriented in such a way that the YY1 protein can position and orient the RNA polymerase II to direct transcription towards the YY motif (**Fig. 4d**; see ref.^42^). In addition, many TF motifs preferentially enriched close to the TSS (**Fig. 4d**). On the 5’ side, the strongest enrichment occurs close to the TSS, slowly decreasing as a function of distance. On the 3’ side, the enrichment declines more sharply so that very little enrichment of most motifs is observed beyond the +20 position from the TSS (**Fig. 4d,e**).

### Prediction of cellular transcriptional activity using the sequence features

To determine how well transcription can be predicted based on the evolved promoter sequences, we used the sequences to train a CNN model (see **Methods**), and to predict the TSS positions genome-wide. To test the CNN, we first used it to score wild-type and mutant forms of the TERT promoter^43,44,45,46^; the model correctly predicted that known cancer-associated mutations^45^ increase the activity of this promoter (**Fig. 4f**; **Fig. S9a-d**). We next used the CNN to predict the positions of active TSSs in GP5d cells. The TSS annotation was derived from the EPD database^47^, and the activity of the TSSs was determined using cap analysis of gene expression (CAGE; see **Methods**). This analysis revealed that promoters evolved from random sequences were more predictive than the genomic sequences themselves (**Fig. 4g**; **Fig. S9e**). A novel mutual information based analysis of interactions learned by the CNN classifiers (see **Methods**) revealed that the classifiers trained on STARR-seq data learned a stronger position-specific signal than the classifiers trained on the EPD data, which relied more on information present at a relatively short region around the TSS (**Fig. S9f,g**; see **Methods**). These results highlight the power of unbiased interrogation of sequence-space that is 100 times longer than the human genome.

### Local enhancer-promoter interactions are additive and non-specific in nature

The binary STARR-seq approach allows identification of interactions between promoters and enhancers. For this analysis, we counted single motif matches at the promoter and enhancer positions, and all pairs of motif matches. When promoters and enhancers were analyzed separately, almost all pairs of TF motifs enriched independently of each other. Strikingly, even across promoters and enhancers, all motifs were independently enriched, suggesting that enhancers activate promoters – but in a very non-specific manner (**Fig. 5a**). Some highly enriched TF-TF pairs, however, displayed weaker activity than that expected from a model that assumes additive action of the enhancer and promoter (**Fig. 5b**). No TF-TF pair was identified that would display substantially stronger transcriptional activity than that expected from independent action of the individual TFs (see also **Fig. S10a**).

**Fig. 5.**
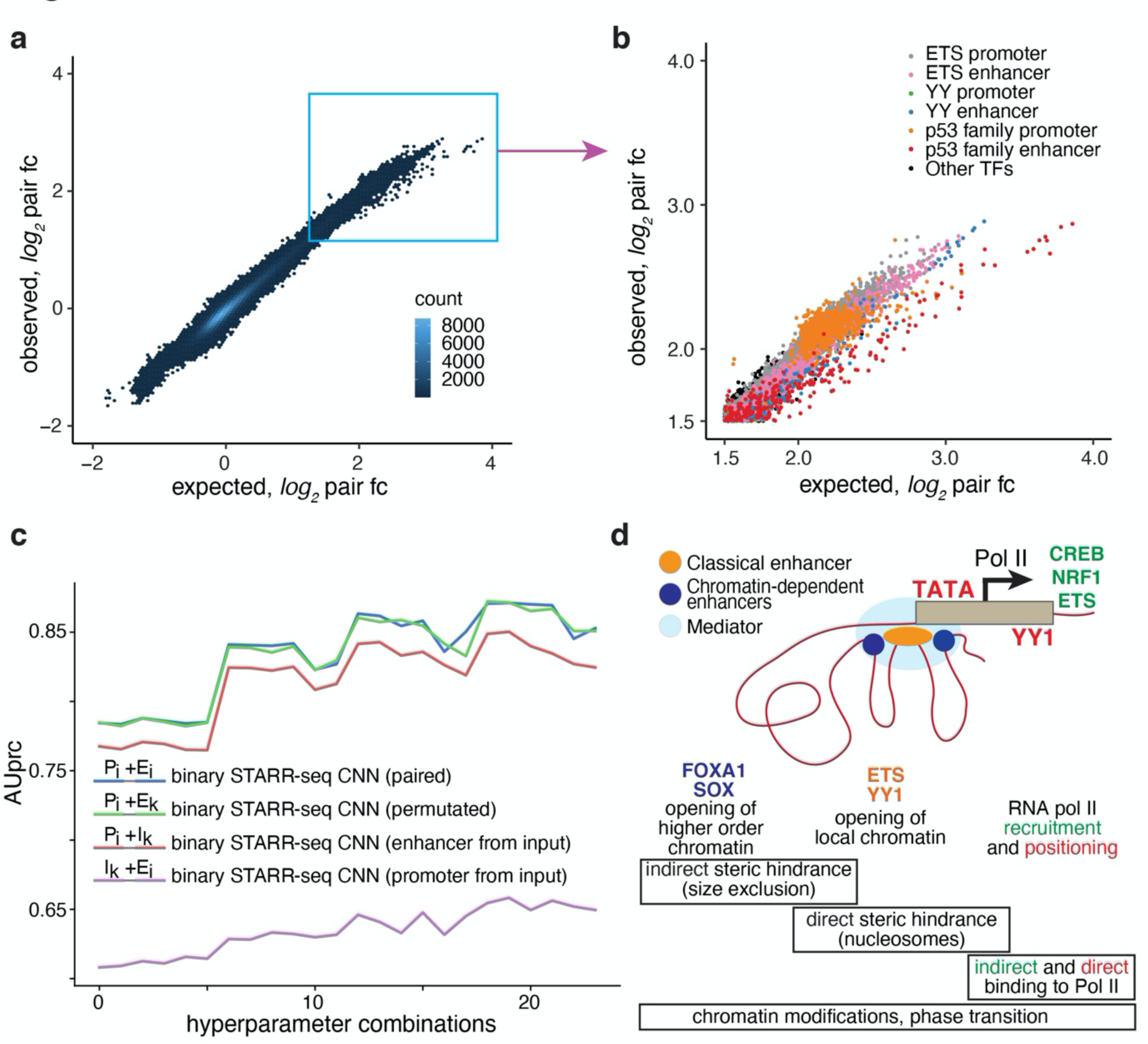
Enhancer-promoter interactions are additive and non-specific in nature. **a,** Plot showing the number of observed promoter-enhancer pairs from the activity measurements from random promoter and random enhancer in the same experiment. For each motif pair, the observed log_2_ fold change of promoter-enhancer pairs compared to input DNA (y-axis) is plotted against the expected change (x-axis) assuming that the promoter motif and the enhancer motif act independently of each other (see Methods for details). The motif matching was done using a p-value threshold 5 × 10^-5^. **b,** Magnified image from the right upper-hand corner of panel (**a**) with the pairs including ETS, p53 family motif and YY1 colored as indicated. **c,** No sequence features that determine specificity of enhancer-promoter interactions can be identified using unbiased machine learning. Four CNN classifiers with identical architecture were trained on different training data sets from the GP5d binary STARR-seq experiment to classify between active and inactive promoter-enhancer pairs. In the “paired” training data the pairing between the promoter and enhancer sequences was retained, whereas in the “permutated” training data, the pairs were shuffled so that any specific interactions between promoter-enhancer pairs are lost. In the “enhancer from input” and “promoter from input” training sets, the promoters and enhancers, respectively, were paired with a randomly sampled sequence from the input library, so the classification is purely based on the promoter or enhancer features. Separate models were trained for 24 different hyperparameter combinations (x-axis; see **Methods** for details): the area under precision-recall curve (AUprc) for all the tested hyperparameter combinations is shown. Note that the classifiers trained on paired data (blue) outperform classifiers trained on enhancer (violet) or promoter (red) data, but not those trained with permutated data (green, paired Student’s t-test p-value ≈1.34x10^-1^). **d,** The relationship between the three TF classes and key processes that control transcription at different scales. TFs affect the gene regulatory process in three ways: First, by directly or indirectly affecting higher-order chromatin structure (left); second, by displacing nucleosomes and opening local chromatin (middle); and third, by recruiting and positioning RNA polymerase II to the gene regulatory elements (right). Gene regulatory unit with classical (orange) and chromatin-dependent (dark blue) enhancers interacting with a gene promoter (brown) is shown. Mediator, a strong feature of both chromatin-dependent enhancers and promoters is shown in light blue. How the hierarchical structure could lead to TFs having chromatin-dependent enhancer (FOXA, SOX), classical enhancing (YY1, ETS), promoting (ETS, CREB, NRF1) and TSS-determining (TATA, YY1) activities is indicated. The relative non-specificity of interactions between TFs, classical enhancers and promoters would be explained by an important role of non-specific molecular interactions such as indirect steric hindrance (size exclusion^72^) and nucleosome-mediated cooperativity (direct steric hindrance^74^) in regulation of transcription. In addition, our general observations are consistent with a role for processes that allow low selectivity such as phase separation and recruitment in the transcription process^75^.

Unbiased machine learning analysis also supported a general mechanism of integration of promoter and enhancer activities (**Fig. 5c**). A classifier using only promoter sequences outperformed a classifier using the enhancer sequences, indicating that the promoter-elements contained more information required for classification of regulatory element activity. As expected, combining the promoters with the correct enhancer sequences increased classification performance significantly. However, permutating the pairings between the promoters and enhancers resulted in similar performance, indicating that there was no predictive power in the specific pairing of individual promoters and enhancers. Taken together, these results indicate that the mechanisms that control transcription are very general, and that the activities of all TFs can independently contribute to transcriptional activity.

## Discussion

Learning the rules by which DNA sequence determines where and when genes are expressed has proven surprisingly hard, despite the availability of full genome sequences of several mammals^48^, extensive maps of genomic features^9, 10, 15, 18^, and genome-scale data about TF protein expression levels and TF DNA binding *in vitro*^6, 8, 49^. Direct comparison of activities between TFs has remained difficult, and therefore we generally lack parameters describing the relative strength of the different sequence features and their interactions that are required to predict their aggregate activity. To address this, we have here defined sequence determinants of human regulatory element activity in an unbiased manner, using a molecular evolution approach where genomic, designed and random sequences are identified that display promoter or enhancer activity.

We found that the cellular gene regulatory system is relatively complex, consisting of several distinct kinds of elements. However, the motif grammar within the elements is relatively loose. In the evolved sequences, precise TF arrangements such as those found in the interferon enhanceosome^50, 51^ are rare, with most elements consisting of TFs acting together largely in an additive manner^30,31,32, 52,53,54,55,56^. Cells are very similar to each other, and the topology of the gene regulatory network is hierarchical, with few TFs displaying very strong transactivation activity. Our findings contrast with the known tissue-specificity of many putative enhancer elements *in vivo*^57, 58^. Interestingly, the level of conservation of many endogenous promoters and enhancers appears to be higher^59^ than the elements selected in our assay (**Fig. S10b)**. The simplest explanation for these two facts is that enhancers *in vivo* evolve to be specific, and that due to the similarity between cells, specificity is more difficult to achieve than activity. Specificity is also required to silence strongly active elements in cell types where a protein is not needed, due to the significant fitness cost of protein expression^60^.

The original functional definition described enhancers as genetic elements that can activate a promoter from a distance, irrespective of their orientation relative to the TSS^1^. We find here that in addition to these elements, two other types of enhancing gene regulatory elements exist: chromatin-dependent enhancers and closed chromatin enhancers (**Fig. 5d**). The chromatin-dependent enhancers are characterized by forkhead motifs, and binding of Mediator and p300 protein, and strong signal for H3K27 acetylation. Unlike classical enhancers, chromatin-dependent enhancers do not transactivate a heterologous promoter strongly, most likely due to lack of binding of TFs with strong transactivator domains. Their presence is, however, strongly predictive of tissue-specific gene expression, suggesting that they act to increase gene expression via chromatin modification or structural changes in higher-order chromatin. Several chromatin-dependent enhancers also combine with a single classical enhancer to form super-enhancers (see **Fig. 2b**), indicating that these elements may be required for driving high levels of gene-expression from distal promoters.

Closed chromatin enhancers, in turn, are located in regions that show little or no signal for DNase I hypersensitivity or ATAC-seq. They are not silenced by CpG methylation. These elements appear to consist of only a single TF (e.g. p53; see also ref.^61^) or a set of closely bound TFs that fit between or associate directly with well-ordered nucleosomes^41^. The prevalence of both the closed chromatin enhancers and chromatin-dependent enhancers suggests that they may contribute significantly to regional control of gene expression^56, 62^.

By using machine learning approaches, we show here that transcriptional activity in human cells can be predicted from sequence features (see also refs^63,64,65,66^). Interestingly, we found that the promoters enriched from completely random synthetic sequences in a single experimental step are even more predictive of transcriptional activity than the genomic sequences themselves. By analysis of *de novo* evolved promoters, we discovered a novel G-rich element that interacts with the TSS, potentially positioning RNA polymerase II to the TSS independently of the TATA box. Overall, TF activities could be classified into three groups: TSS position determining activity (e.g. TATA-box, YY), short range promoting activity (e.g. NRF1), and enhancing activity (many TFs). We did not detect a separate class of distal enhancing activity, suggesting that activities that would allow an enhancer to selectively act at a very long range is likely to be associated with chromatin-dependent enhancers and not classical enhancers^67,68,69,70,71^. The three classes of activities detected are not mutually exclusive, suggesting that TFs act at multiple levels and/or scales to regulate transcription (**Fig. 5d**). For example, YY1 acts both as an enhancing TF and as a TSS-determining one, and that FOXA motifs are present at both chromatin-dependent and classical enhancer elements. Our results thus indicate that gene regulatory elements are not the atomic units of gene expression, and that TF motifs should ultimately be the basis of analysis and prediction of genomic gene regulatory activity.

Our random promoter-enhancer design allowed unbiased discovery of features that facilitate interactions between classical enhancers and promoters at a relatively short range. No specific pair of motifs controlling such interactions was found. This, together with the fact that no specific TF that only acts from an enhancer was found is consistent with a generic and indirect mechanism of action, where the activities of individual TFs bound to an enhancer are aggregated, and their total activity then activates the promoter. Molecularly, these results are consistent with mediation of the effect by the least specific type of biochemical interaction, steric hindrance. The simplest mechanism for enhancer action would involve opening of higher order and local chromatin in such a way that the steric hindrance that prevents large macromolecular complexes such as mediator or RNA polymerase II from loading to DNA is decreased (**Fig. 5d**; see ref.^72^). In summary, we show here that direct experimentation to interrogate transcriptional activities of sequences that are on aggregate >100 times longer than the human genome can be used to determine mechanisms of action of, and interaction between, gene regulatory elements. The experiments revealed unexpected simplicity of gene regulatory logic. The discovery of the relative simplicity of the interactions, together with the ability to measure transcriptional activities of all TFs in a cell represents a significant step toward achieving the ultimate aim of regulatory genomics – predicting gene expression from sequence.

## Methods

### STARR-seq vector design

We designed a modified STARR-seq reporter construct pGL4.10-Sasaki-SS (a) based on the earlier published design^21^ in pGL4.10 backbone (Promega, #E6651). The sequence between SacI and AfeI was replaced with a sequence containing the chicken lens δ1-crystallin gene (Sasaki) promoter^76^, a synthetic intron (pIRESpuro3, Clonetech, #631619), an ORF (fusion of Nanoluc-EmGFP), homology arms for library cloning with AgeI and SalI RE sites flanking the ccdB gene, a small 52 bp DNA stuffer (a part of the neomycin resistance cassette) and a 20 bp sequence from the 3’-Illumina adapter for optimally sized final library for Illumina sequencing, and the SV40 late polyA-signal from pGL3 backbone (Promega, #E1751).

To enable the analysis of CpG methylation on enhancer activity, we designed modified STARR-seq vectors in a CpG-free backbone with Lucia reporter gene (Invivogen, #pcpgf-promlc) driven either by the EF1α promoter (b. pCpG-free-EF1α-SS) or the Sasaki promoter (c. pCpG-free-Sasaki-SS-v1) as above. To facilitate the cloning of the synthetic DNA library to the 3’-UTR of the reporter gene, the cloning cassette from the pGL4.10-Sasaki-SS vector (a) containing the homology arms with AgeI and SalI RE sites, the 52 bp DNA stuffer, and the 20 bp sequence from the 3’-Illumina adapter as above was introduced to the CpG-free vectors using the NheI site.

Standard Illumina adapters harbor CG dinucleotides, and to make our modified STARR-seq design completely CpG-free, we designed custom adapters for Illumina sequencing (see Oligos 3 and 4 in **Table S2**). To accommodate the cloning of genomic DNA and random sequence inserts with flanking CpG-free custom adapters, the cloning cassette in CpG-free-Sasaki-SS-v1 was modified by removing the 3’-Illumina adapter and the 52 bp stuffer. In addition, this vector was further improved by replacing the AgeI and SalI RE sites with the AflII and PvuII sites devoid of CG dinucleotides, and by introducing a DNA stuffer of 1.2 kb between the RE sites to the resulting pCpG-free-Sasaki-SS-v2 vector (d) to unambiguously detect and purify the linearized reporter backbone for downstream cloning.

For the binary STARR-seq approach in which random sequences were cloned as both promoters and enhancers, the pCpG-free-Sasaki-SS-v2 vector (d) was modified by replacing the Sasaki promoter with a custom CpG-free 5’-adapter sequence and an AgeI RE site, and by introducing a SalI RE site and a custom CpG-free 3’adapter immediately downstream of the ORF. Moreover, to optimize the random promoter and random enhancer library size for Illumina sequencing, the Lucia reporter gene was replaced by a small eleven amino acid ORF from *Drosophila melanogaster* (Dm tal-1A) in the pCpG-free-promoter-enhancer-SS vector (e). The cloned random promoter-random enhancer input library is paired-end sequenced to map the promoter-enhancer pairs, and thus the random enhancer sequences obtained after sequencing the reporter-specific RNA library can be used to identify the corresponding promoter sequence from the input library. In total, the constant sequence between promoter and enhancer elements is 872 bp in the pCpG-free-Sasaki-SS-v2 construct and 215 bp in the pCpG-free-promoter-enhancer-SS construct.

The new reporter vectors (a-e) were used in different experiments as summarized below, and their complete sequences are provided in **Table S1**.

**a.** pGL4.10-Sasaki-SS (5,754 bp) used for experiments with synthetic motif library shown in Fig. S2b,c
**b.** pCpG-free-EF1α-SS (3,497 bp) used for experiments with synthetic motif library shown in **Fig. 1b-e, Fig. S2a,b,d**
**c.** pCpG-free-Sasaki-SS-v1 (3,388 bp) intermediate plasmid not used in the experiments
**d.** pCpG-free-Sasaki-SS-v2 (4,458 bp) used for all experiments with genomic fragments and random enhancer (N170) sequences
**e.** pCpG-free-promoter-enhancer-SS (2,551 bp) used for all experiments with random promoter (N150)-random enhancer (N150) sequences

### STARR-seq reporter library construction and cloning

We have generated STARR-seq reporter libraries from rationally designed oligonucleotides harboring TF binding motifs, from fragmented human genomic DNA, and from random oligonucleotide sequences as detailed below. All the oligonucleotides that were used for the cloning of the libraries were purchased from Integrated DNA Technologies and their sequences are provided in **Table S2**.

### TF motif input DNA library

A pool of 92,918 oligos with a length of 79 nucleotides (nt) was designed with a 49-nt variable region and two 15-nt flanking regions with constant sequences for the library cloning (see Motif library design for more details), and synthesized by CustomArray Inc. The oligo pool was prepared for cloning in a two-step protocol using Phusion DNA polymerase (Thermo Fisher) and Oligos 1 and 2. Frist, 2.5 pmol of the oligo pool was double-stranded using 100 pmol of Oligo 2 in two parallel reactions [98 C for 3 minutes (min), followed by 5 cycles of 98 C for 10 seconds (s), 55 C for 15 s, 72 C for 15 s, and the final extension at 72 C for 2 min]. The two reactions were then split into ten reactions and after adding 10 pmol of the Oligo 1 to the reactions the PCR was performed for ten additional cycles using the same conditions. Ten PCR reactions were pooled and the 127-bp product was gel-purified. The pGL4.10-Sasaki-SS (a) and pCpG-free-EF1α-SS (b) vectors were linearized by digestion with AgeI and SalI for 3 h at 37 C and gel-purified. For each backbone, the In-Fusion cloning (Clonetech) was performed in 20 reactions using 200 ng of linearized vector and 50 ng of double-stranded oligo pool according to the manufacturer’s instructions. Five In-Fusion reactions were pooled and purified using MinElute PCR purification columns (Qiagen) and eluted in 12.5 ul nuclease-free water per column. The bacterial transformation was done by electroporation using Gene Pulser Xcell (Biorad) in 20 parallel reactions with 2.5 ul of purified eluate and 20 ul of E. cloni 10G SUPREME (Lucigen) or Transformax EC100D pir-116 (Lucigen) electrocompetent cells for the pGL4.10-Sasaki-SS (a) and pCpG-free-EF1α-SS (b) vectors, respectively, using the manufacturer’s recommendations. To each electroporation cuvette, 1 ml of recovery media (Lucigen) was added, and the cells were incubated for 1 h at 37 C and 250 rpm. All cultures were then pooled and added to 5L of LB media with 100 ug/ml ampicillin (pGL4.10-Sasaki-SS; a) or 25 ug/ml zeocin (pCpG-free-EF1α-SS; b) and grown overnight at 37 C and 250 rpm until the O.D. reached 1.0. The bacterial cells were harvested by centrifugation and plasmid DNA was isolated using EndoFree Plasmid Giga kit (Qiagen).

### Genomic DNA input library

Genomic DNA was isolated from GP5d colon cancer cells using DNeasy Blood and Tissue kit (Qiagen) and treated with RNAseA (Thermo Fisher) followed by purification. The genomic DNA was fragmented to an average size of ∼500 bp using Covaris S220 according to the manufacturer’s recommendations. The fragmented genomic DNA (400 ng per reaction in ten reactions) was end-repaired, dA-tailed, and ligated to custom CpG-free annealed adapters (Oligo 3 - Custom CpG-free P7 adapter and Oligo 4 - Custom CpG-free P5 adapter; annealed according to the standard Illumina protocol). The adapter-ligated gDNA was purified and amplified in 20 reactions using KAPA HiFi master mix (Roche) and Oligos 5 and 6 which add the homology flanks for the NEB HiFi DNA assembly. The PCR product was pooled and purified using 0.8x volume of AMPure XP beads (Beckman Coulter) using the manufacturer’s instructions followed by MinElute column purification. The pCpG-free-Sasaki-SS-v2 vector (d) was linearized using AflII and PvuII for 3h at 37 C and gel-purified. The PCR fragments were recombined to the vector in 50 parallel NEBuilder HiFi DNA assembly reactions (NEB) according to the manufacturer’s instructions using 150 ng of linearized vector and 50 ng of custom CpG-free adapter-ligated genomic DNA. The reaction products were pooled (five reactions per column) and purified using MinElute columns (Qiagen) and eluted in water. The bacterial transformation was done in 50 reactions using 2.5 ul of purified eluate and 20 ul of Transformax EC100D pir-116 (Lucigen) by electroporation (Gene Pulser Xcell, Biorad) using the manufacturer’s recommendations. To each electroporation cuvette, 1 ml of recovery media (Lucigen) was added, and the cells were incubated for 1 h at 37 C and 250 rpm. The 50 cultures were then pooled and added to 6L of LB media with 25 ug/ml zeocin and grown overnight at 37 C and 250 rpm until O.D. reached 1.0. The bacterial cells were harvested by centrifugation and plasmid DNA was isolated using EndoFree Plasmid Giga kit (Qiagen).

### Random enhancer oligonucleotide DNA input library

The random enhancer library was constructed from a 200-nt single-stranded Ultramer oligonucleotide (Oligo 7) harboring a 170-nt random sequence (170N) flanked by 15-nt constant sequences for library cloning. Double-stranded library was produced by employing a similar PCR strategy as described above for the TF motif library using Phusion DNA polymerase (Thermo Fisher) and Oligos 8 and 9 that introduce custom CpG-free sequencing adapters and flanking sequences homologous to the pCpG-free-Sasaki-SS-v2 vector (d). The NEB HiFi assembly between the random enhancer PCR product and the AflII-PvuII fragment from the pCpG-free-Sasaki-SS-v2, as well as electro-transformation and plasmid DNA isolation were performed as described above for the genomic DNA library.

### Random promoter and random enhancer oligonucleotide DNA input library

The random promoter-random enhancer library was constructed by using two 190-nt single-stranded Ultramer oligonucleotides (Oligos 10 and 11) with 150-nt random sequences (150N). Each oligo harbors two 20-nt constant sequences that facilitate the library cloning to the pCpG-free-promoter-enhancer-SS vector (e). First, the constant sequences at the 3’ ends of the oligos anneal to the pCpG-free-promoter-enhancer-SS vector in a PCR reaction with Phusion DNA Polymerase (Thermo Fisher), amplifying the region between AgeI and SalI sites from the backbone. Then, the PCR product constituting the random promoter-random enhancer library (of size 555 bp) was cloned into the pCpG-free-promoter-enhancer-SS vector (e) linearized using AgeI-SalI using the constant sequences introduced by the 5’ends of the oligos. The NEB HiFi assembly, electro-transformation, and plasmid DNA isolation were performed as described above for the genomic DNA library.

### CpG methylation of STARR-seq input DNA library

The genomic DNA library cloned into the pCpG-free-Sasaki-SS-v2 vector was methylated using bacterial CpG methylase *M.SssI* (NEB). The reaction was performed for 4 h at 37 C as per manufacturer’s recommendation with the reaction volumes scaled for 62.5 ug of plasmid DNA per reaction, and inactivated for 20 min at 65 C, followed by purification and ethanol precipitation to have the transfection-ready methylated library.

### Cell lines and generation of TP53-null cell line by genome editing

The cell lines used in this study were colon cancer cell line GP5d (Sigma #95090715), liver cancer cell line HepG2 (ATCC #HB-8065), and retinal pigmented epithelial cell line hTERT-RPE1 (ATCC #CRL-4000). The cells were maintained in their respective media (GP5d in DMEM, HepG2 in MEM and RPE1 in DMEM/F12) supplemented with 10% FBS, 2nM L-glutamine, and 1% Penicillin-Streptomycin.

*TP53*-null GP5d cell line was generated by targeting exon 4 of the *TP53* gene using CRISPR-Cas9 genome editing. sgRNA duplex was annealed from crRNA (Oligo 12) and tracrRNA with atto550 (Integrated DNA Technologies) and used for ribonucleoprotein (RNP) complex formation with Cas9-HiFi protein (Integrated DNA Technologies) as per manufacturer’s recommendations, and the RNP complex was transfected to GP5 cells using CRISPRMAX reagent (Invitrogen). On the next day, atto550-positive cells were FACS sorted and single cell colonies were cultured to produce a clonal *TP53*-null cell line. The clonal cells lines were screened for TP53 depletion by western blotting and the clones were verified by Sanger sequencing of PCR products from isolated genomic DNA using Oligos 13 and 14.

### Transfection and RNA isolation

In the STARR-seq experiments, 1 ug of each input library DNA was transfected per million cells. For TF motif DNA libraries, a total of 50 and 35 million GP5d cells were transfected for the libraries in pGL4.10-Sasaki-SS (a) and pCpG-free-EF1α-SS (b) vectors, respectively. Experiments were performed in two replicates with random enhancer libraries in GP5d and HepG2 cells and with random promoter-enhancer libraries in GP5d cells (250 million cells per each replicate). Genomic STARR-seq experiments were performed in two replicates in HepG2 cells (170 million cells per replicate) and in four different conditions in GP5d cells (wild type and TP53-null GP5d cells using both methylated and non-methylated input DNA libraries, 500 million cells per condition). For random promoter-enhancer libraries in HepG2 and RPE1 cells, a total of 400 and 480 million cells were transfected, respectively. Briefly, a day before transfection, 6.7-10 million cells were plated per 15-cm dish in their respective media without antibiotics. Next morning, plasmid DNA was mixed with transfection reagent optimized for each cell line [Transfex (ATCC) for GP5d, Transfectin (BioRad) for HepG2, and FuGENE HD (Promega) for RPE1] in 1:3 ratio in Opti-MEM media (Gibco), incubated for 15 min at RT, and added dropwise to the cells. The cells were incubated for 24 h in a 37 °C incubator with 5% CO2.

Cells were harvested and total RNA isolated 24 h after transfection using RNeasy Maxi kit (Qiagen) with on-column DNaseI digestion as per manufacturer’s instruction. PolyA(+) RNA fraction was purified using Dynabeads™ mRNA DIRECT™ Purification Kit (Invitrogen, #61012) as per the manufacturer’s recommendation followed by DNase treatment using TurboDNase (Ambion) and purification using RNeasy Minelute kit (Qiagen) as described previously described^21^.

### STARR-seq reporter library and input DNA library construction

The reporter library preparation protocol was adapted from ref.^21^ essentially in all steps but with primers matching our modified STARR-seq vectors. First strand cDNA synthesis was done with 2.5-5 ug of polyA(+) RNA and with Superscript III (Invitrogen, #18080-044) using a reporter-RNA specific primer (Oligo 15) in 10-20 reactions depending on polyA(+) RNA yield. This was followed by RNase A treatment for 1 h at 37°C and purification using MinElute PCR purification columns (Qiagen). cDNA amplification was performed with reporter-specific nested cDNA primers (Oligos 16 and 17 for libraries in vectors d and e, Oligos 16 and 18 for libraries in vector b, and Oligos 19 and 18 for libraries in vector a) using KAPA HiFi PCR Master mix (Roche) in the same number of reactions as done for the reverse transcription (98 C for 2 min, followed by 15 cycles of 98 C for 15 s, 65 C for 30 s and 72 C for 30-70 s). The PCR products were purified using 0.9X AMPure XP beads as per manufacturer’s instruction followed by elution in nuclease-free water. The final PCR reactions to produce Illumina-compatible sequencing libraries were prepared from the entire amplified cDNA using KAPA HiFi PCR Master mix (Roche) at 98 C for 2 min, followed by 8-10 cycles of 98 C for 15 s, 65 C for 30 s and 72 C for 30 s. The primers used for different libraries are as follows: TF motif libraries in pGL4.10-Sasaki-SS (a) and pCpG-free-EF1α-SS (b) vectors were amplified using standard Illumina Universal and index primers (NEB #E7335S) and sequenced using standard Illumina chemistry. Genomic DNA and random enhancer libraries in pCpG-free-Sasaki-SS-v2 vector (d) were amplified using custom CpG-free primers (Oligos 20 and 21) and random promoter-random enhancer libraries in pCpG-free-promoter-enhancer-SS vector (e) using Oligos 22 and 21. All custom CpG-free libraries were sequenced using custom read 1 and read 2 primers (Oligos 23 and 24) and custom i7 index read primer (Oligo 25). For preparing sequencing libraries from the input DNA, plasmid DNA from each library design was amplified in ten parallel reactions (10 ng DNA per reaction) using Phusion DNA Polymerase (Thermo Fisher) as above. For TF motif libraries in pGL4.10-Sasaki-SS (a) and pCpG-free-EF1α-SS (b) vectors, standard Illumina primers (NEB #E7335S) and sequencing chemistry were used, and for all the libraries in pCpG-free-Sasaki-SS-v2 (d) and pCpG-free-promoter-enhancer-SS (e) vectors, Oligos 20 and 21 were used for PCR amplification and Oligos 23-25 for sequencing. All libraries were sequenced either single-end or paired-end as per Illumina’s standard instructions and protocols on suitable Illumina platforms like MiSeq, NextSeq500, HiSeq4000 and NovaSeq.

### Template switch library preparation

For generating a sequencing library using a template switch strategy, a 40-ug aliquot of total RNA from the random promoter-random enhancer STARR-seq experiment in GP5d cell line was used. Briefly, TurboDNase-treated RNA was incubated at 72 C for 3 min with a custom biotinylated STARR-seq specific RT-primer (Oligo 26) and dNTPs in two reactions having 25 ng of RNA per reaction, followed by immediate chill on ice. The first strand synthesis was performed using 100 units of Superscript IV RT (Invitrogen) in 1x SS-IV RT buffer containing 10 units of RNase OUT (Invitrogen), 5 mM DTT, 6 mM MgCl_2_ (Sigma), 1M Betaine (Sigma) and 1 uM custom template switch oligo (Oligo 27) compatible with the custom CpG-free Illumina sequencing by incubating the reactions at 50 C for 15 min followed by 80 C for 10 min. cDNA was purified using AMPure XP beads (Beckman Coulter) and one cDNA reaction product was split into two for PCR amplification using KAPA HiFi master mix (Roche) together with USER enzyme (NEB) and Oligos 28 and 17 for enrichment of STARR-seq reporter-specific template (98 C for 3 min, followed by 15 cycles of 98 C for 20 s, 67 C for 15 s, 72 C for 6 min and final extension at 72 C for 5 min). The PCR product was purified using AMPure XP beads (Beckman Coulter) followed by a second PCR for a total of 5 cycles for Illumina library preparation using custom CpG-free Oligos 20 and 21. The final library was purified and sequenced using Oligos 23-25 on NextSeq and NovaSeq platforms.

### Chromatin immunoprecipitation (ChIP-seq) and gene expression analysis (RNA-seq and CAGE)

Chromatin immunoprecipitation (ChIP) was performed as previously described^8^. For analyzing genomic occupancy of TP53, cells were treated with 350 uM 5-fluorouracil (Sigma) 24 h before harvesting the cells. Briefly, fresh formaldehyde-crosslinked chromatin from GP5d cells was used to immunoprecipitate DNA using Dynal-bead coupled antibodies for H3K27ac, H3K9me3, and H3K27me3 (C15410196, C15410193, and C15410195, Diagenode, respectively), FOXA1 (ab23738, Abcam), p53, HNF4a, and CTCF (sc-135773x, sc-8987x, and sc-15914x, Santa Cruz, respectively), SMC1 (A300-055A, Bethyl lab), and for normal rabbit, mouse and goat IgG (sc-2027, sc-2025, and sc-2028, Santa Cruz, respectively), followed by standard ChIP-seq library preparation for Illumina sequencing. The libraries were single-read sequenced on HiSeq4000. The Illumina raw files were demultiplexed using bcl2fastq conversion, followed by alignment to the human genome (hg19) using bowtie2^77^ and peak calling (narrow peaks for TF ChIP-seq and broad peaks for histone modifications) was performed using MACS2^78^ using default parameters. Super-enhancers were calculated using SMC1 and H3K27ac ChIP-seq data using the ROSE pipeline^79^. The peak files were filtered for the ENCODE blacklisted region (accession ENCSR636HFF) before further downstream analysis.

For gene expression analysis, the transcript abundances (transcripts per million; tpm) for three replicate samples were taken from RNA-seq data from previous studies for GP5d (from ref.^80^) and HepG2 (from ref.^81^). The expression of each TF in GP5d cells was summarized by taking the mean expression of its most highly expressed transcript over the replicates. For comparison between GP5d and HepG2, pseudocount 1 was added to the expression values (tpm), and the genes with mean tpm < 2 across all experiments were excluded (36% of the protein coding genes). Differential expression between GP5d and HepG2 cell lines was then estimated for the remaining 12,586 genes with the limma package^82^ using eBayes with parameter trend set to true. CAGE library was prepared from total RNA isolated from GP5d cells as described in ref.^14^ with an input of 1 µg total RNA. Sequencing of the CAGE library was carried out on HiSeq 2000 (Illumina).

### Chromatin accessibility (ATAC-seq)

The ATAC-seq library was prepared from 50,000 GP5d cells as previously described^83^. The cells were washed in ice-cold PBS and resuspended in 50 ul of lysis buffer and incubated for 10 min on ice. The pellet from lysed was transposed with Tn5 transposase in 2X tagmentation buffer (Illumina kit) and incubated for 30 min at 37 C. The reaction was purified using a MinElute purification kit and eluted in nuclease-free water. The samples were amplified for 5-8 cycles as determined by qPCR for Illumina sequencing using Nextera library preparation kit (Illumina) and samples were paired-end sequenced on HiSeq4000. The samples were demultiplexed and paired-end fastq files were processed using an in-house pipeline comprising of TrimGalore (https://www.bioinformatics.babraham.ac.uk/projects/trim_galore/), BWA aligner^84^, Picard (http://broadinstitute.github.io/picard/) and broad-peak calling by MACS2^78^. The peak files were filtered for the ENCODE blacklisted regions as described earlier.

### Active transcription factor identification (ATI) assay

The assay was performed *in vitro* by mixing 5 μl nuclear protein extracted from GP5d cells (2ug/ul), 5 μl 140 bp double stranded DNA (dsDNA) oligos^29^ containing 40 bp random sequence in the middle (10 pmol), and 5 μl 3 × protein binding buffer (420 mM KCl, 15 mM NaCl, 3 mM K_2_HPO_4_, 6 mM MgSO_4_, 300 μM EGTA and 9 μM ZnSO_4_, 60 mM HEPES, pH = 7.5) and incubating for 30 min at room temperature. The poly-dIdC was supplemented in the reaction (5 ng/ μl final concentration) to decrease non-specific binding. Electrophoretic mobility shift assay (EMSA) was then conducted using commercial DNA Retardation Gel (Invitrogen, #EC63652BOX) in 0.5 × TBE buffer (1 mM EDTA in 45 mM Tris-borate, pH 8.0) at 106 V voltage for 70 min. The gel above the 300 bp DNA marker was collected, eluted in 300 μl Tris buffer (10 mM Tris-Cl, pH 8.0) and incubated at 65 °C for 3 h. The eluted DNA was amplified with Phusion polymerases (Thermo Scientific, #F530L); 4 pmol of each primer were used for the amplification. Before the final step of amplification, the same amount of primers was added to convert the remaining single stranded DNA (ssDNA) to dsDNA. The amplified DNA library was incubated again with an aliquot of the same protein extract as above and the whole process was repeated for three more times. The PCR products from different cycles of ATI were purified and sequenced by Illumina Hiseq4000.

### Transient transcriptome sequencing (TT-seq)

Transcribed enhancer regions defined using TT-seq data are based on Lidschreiber et al., 2020 (manuscript in preparation). Briefly, TT-seq from GP5d colon cancer cells was performed in two biological replicates as described^85^. TT-seq libraries were sequenced to a depth of ∼120 million uniquely mapped paired-end reads and data analysis for identification of genomic intervals corresponding to continuous uninterrupted transcription (defined as transcription unit, TU) was performed using GenoSTAN^86^. TUs were classified into two groups: those overlapping with annotated protein-coding genes were defined as mRNAs and remaining as non-coding RNAs (ncRNAs). Putative enhancer RNAs (eRNAs) were further subclassified using histone modification patterns by defining a large set of putative enhancer regions from the publicly available datasets covering the non-coding regulatory genome^9, 57^. eRNAs were further defined from the pool of ncRNAs on the basis of three criteria; first, origin should overlap within TSS ±500 bp (enhancer region), second, it should be outside of TSS ± 1 kbp (promoter region), and third, transcription is bidirectional as measured with TT-seq. This annotation led to identification of 6774 enhancer regions in GP5d colon cancer cells. The annotated enhancers were filtered for blacklisted regions as described before and also for chromosomes other than 1-22 and X.

### Motif collection

For testing activities of known TF motifs, a set of 3226 HT-SELEX motifs were collected (Refs ^6, 8, 87^ and unpublished draft motifs). A more compact set of motifs representing different binding specificities was generated by first constructing a dominating set (880 PWMs) covering motifs from the above sources using the same method and motif distance threshold as in ref.^6^. Then, in order to retain information of TF binding differences between methylated and non-methylated DNA ligand^8^, for each methyl (or non-methyl) motif in the dominating set, the closest non-methyl (methyl) motif of the same TF was added to the representative set (respectively) if it was not yet in the set. This resulted in a representative set of 1121 HT-SELEX motifs (**Table S3**). In cases where HT-SELEX motifs for several TFs are highly similar, the motifs have been named in figures according to TF class or subclass, in order to highlight that we do not know which of the TFs that have similar motifs binds to the motif in the cells. Same principle has been applied also when specificities of closely related TFs have not been measured and thus can reasonably be expected to be similar. **Table S5** shows the naming for the motifs in each figure. A control set of reversed but not complemented motifs was generated by reversing the column order of each motif matrix. Additionally, the following promoter core motifs were collected for TSS analyses from literature: TATA box, Initiator, CCAAT-box, GC-box from ref.^88^, and BRE, MTE, DPE from ref.^89^.

### Motif library design

The synthetic oligo library design contained the 1121 HT-SELEX motifs in various sequence patterns. A pattern is defined as the combination and number of the motif consensus sequences, their relative orientation and spacings, and positions of degenerate N bases. Each of 727 monomeric and homodimeric motifs was included in the following patterns: consensus and its reverse complement (if palindromic added twice), reversed but not complemented control, each position of consensus at a time replaced with a degenerate base N, and two and three copies of the consensus sequence in different orientations and spacings (three copies only for the motifs shorter than fifteen bases). Putting a degenerate base N at every position (in total 10,041 positions) generated in total 30,123 mutant consensus sequences.

In the patterns containing two and three copies of the consensus, the most defined position of the motif (position with maximum probability for any base in any position) was replaced with N. The two copy patterns included three relative orientations of the consensus C and its reverse complement R (CC, CR, and RC) and three copy patterns included four relative orientations (CCC, CCR, CRC, RCC). In the case of two copies, each orientation was included with all gap lengths from zero to six bases between the two copies. In the case of three copies the gap length was varied from zero to four bases (oligo length permitting) but the same gap length was used for both gaps in one pattern. Also, the consensus and reverse complement of the 394 heterodimeric motifs^36^ was included. A subset of 245 heterodimeric motifs were also cut into two half-sites and the half-sites were added in the same patterns as two copies of a monomeric motif.

Each of the 43,251 motif patterns was embedded in two different sequence contexts. The contexts were chosen from two human genomic loci (context 1, chr10:77103489-77103535 and context 2, chr8:21525556-21525602, in hg19 coordinates) that do not contain high affinity sites of known motifs. Finally, 2-6 bases long random sequence (UMI) was put to the 5’ end of the sequence to create an approximately uniform base distribution for sequencing. Both context sequences were also included alone with five different UMI lengths. Thus, in total 86,512 sequences were used for testing TF activity. The sequences and their embedded motif patterns are given in **Table S4**. The remaining 6,406 oligos from the total 92,918 sequence patterns were composed of 3,576 SELEX nucleosome bound and unbound sequences, 1,182 draft HT-SELEX models for RNA binding proteins^90^ and 1,648 for tiling genomic regions corresponding to enhancer regions for MYC and CCND2 (**Table S4**).

### Motif library complexity

Based on the sequencing of the pCpG-free-EF1α-SS input library, the motif library was estimated to contain approximately 26.9 × 10^6^ distinct sequences when taking into account different UMIs (corresponding in total 1.3x10^9^ bp), read counts are given in **Table S7**.

### Enhancer activity of TF motif consensus sequences

For each TF consensus pattern, the reads containing the pattern were counted separately for the two sequence context and counts less than five in RNA and input DNA together were discarded from further analysis. The fold change between RNA and input DNA was estimated using the function PsiLFC in R package lfc version 0.2.1^91^ for each pattern in each context. To summarize the activity of one, two, or three copies of the motif, the median fold change of all the patterns in both contexts containing the given number of the consensus sequences was used. For an individual consensus sequence, this included the patterns containing it or its reverse complement in both contexts. In the case of two or three copies, all consensus spacings and orientations were summarized together. The average fold changes over all motifs in one, two, three copies were only calculated from those that could be detected with all copy numbers. If several sites acted without synergy, we assumed log_2_ fold changes to grow linearly as a function of number of sites. The motifs representing heterodimers were excluded from the analysis.

### Analysis allowing base substitutions and generation of activity position weight matrix

For each pattern in a sequence context, the number of matching reads was counted both for the consensus sequence and its variations containing the designed base substitutions. The patterns detected at least 100 times in each context in RNA and input DNA together were considered. The fold change of a motif pattern in a context was estimated as the ratio of RNA and input DNA counts (with pseudocount one) normalized using the total count of all motif patterns considered. The activity PWM was generated using the counts of the consensus sequence and all its single base substitutions. For each position of the consensus, the number of times each base was observed at that position in the sequences otherwise matching the consensus was counted both in RNA and input DNA. Finally, the PWM was constructed by dividing the RNA count by input DNA count in each position (adding pseudocount one).

### Preprocessing of random enhancer-library sequences

First, 150 bases long STARR-seq RNA and input DNA paired-end reads were combined using the FLASH program^92^ and only combined sequences of length 170 were chosen. Duplicate reads were removed by sorting the sequences four times based on 45 bases long non-overlapping subsequences from base 6 to 165 and taking only one sequence per identical subsequence at each sort step. This ensured that from sequences that had Hamming distance less than 4, only one was taken. Only one representative sequence from the similar sequences was used for downstream analysis, so each sequence is either present or absent in the sample. The sequences are sampled from a huge input DNA library which prohibits precise determination of initial input frequencies of individual sequences. Thus, our analysis relies on finding common features of different selected sequences instead of their counts. The resulting numbers of preprocessed sequences used in downstream analysis are shown in **Table S7**.

### Genomic library complexity analysis

Genomic STARR-seq input DNA library complexity was estimated using the preseq program^93^ lc_extrap tool that estimates how many distinct fragments would be observed based on reads in an initial sample if a given number of reads was sequenced. The preseq lc_extrap tool was given the mapped paired-end reads (bam) as input and the expected yield of 2.09 × 10^9^ distinct fragments (assuming 2 × 10^10^ reads were sequenced) was used as the complexity estimate (10^12^ bp). Thus, the library was estimated to cover the human genome (3.2 × 10^9^ bp) with 1.53 bp resolution on average. The resolution of STARR-seq output fragments in highly active genomic regions was broadly consistent with the high input resolution, taking into account that the fragment activity depends on which part of the regulatory element it covers.

### Random library complexity and information analysis

The complexity of random enhancer STARR-seq input DNA library was estimated by first creating a robust set of sequences originating from the same clones and their counts. This was done by clustering a randomly sampled set of input sequences using starcode^94^ so that first reads which edit distance four or less were connected and then reads in the same connected component were put to one cluster. The sizes of the sequence clusters were then given as input to preseq resulting in estimated 2.4 × 10^9^ distinct sequences (corresponding in total 4.1 × 10^11^ bp). The same approach gave an estimate of 0.9 × 10^9^ promoter sequences (1.4 × 10^11^ bp) and × 10^9^ enhancer sequences (1.8 × 10^11^ bp) in the binary STARR-seq input library. To confirm that these extremely complex libraries are effectively transfected to the cells, we estimated the number of plasmid copies per cell based on the comparison of read coverage between plasmid and genomic DNA from a control ChIP-seq experiment using a non-specific IgG antibody. This analysis revealed over 2500 plasmid copies per cell and based on this we estimate that each distinct random enhancer sequence was transfected to cells over 500 times on average.

To compare to the genomic conservation, we assumed that a TF motif typically has ∼15 bits of information content. As it can be placed in ∼320 positions in a 170 bp long random enhancer, one TF binding site contributes approximately 15-log_2_(320) = 6.7 bits of information (see ref.^95^). Thus, a site would correspond to approximately 7 conserved bases (>1 bit of information).

### Genomic STARR-seq analysis

First, demultiplexed Illumina STARR-seq RNA and input DNA paired-end reads (trimmed to common length 2 x 37 bp) were aligned to the human genome (hg19) using bowtie2^77^. Before peak calling the mapped read pairs were deduplicated and paired-end reads that had mapping quality <20 or mapped in a discordant orientation were discarded. MACS2^78^ was used to call peaks in paired-end mode (-f BAMBE) so that the fragment endpoints were inferred from alignment results. Input DNA was used as control in peak calling and the called peaks were filtered for the ENCODE blacklisted regions (accession ENCSR636HFF) before subsequent downstream analysis. For masking repeats, a RepeatMasker file from UCSC table browser (for hg19) was used.

All STARR-seq RNA fragments from each cell line were used in peak calling (**Table S7**) unless otherwise stated. Peaks were also called separately from two replicates in HepG2 cell line with 3295 peaks overlapping out of 6414 and 7376 peaks. IDR software^96^ was then used together with combined sample peaks to call high confidence peaks (2186 peaks with IDR < 0.1). In GP5d cells, genomic STARR-seq analysis was performed in two sublines (p53 wt and null) under two conditions (methylated or not), but due to the very large size of the experiments, replicates were not included for each condition. To enable calling reproducible peaks from the GP5d data, we utilized an internal control approach wherein two sets of peaks are built from a single replicate by splitting the fragments to two sets based on their mapping to either even or odd positions of the genome. These “in silico” replicates were then used for IDR analysis resulting in 1970 and 3250 high confidence in silico peaks (IDR < 0.1) in HepG2 and GP5d wt cells, respectively. Comparison of the high confidence peaks from biological and in silico replicates in HepG2 cells revealed that this IDR method yields similar peak-calls (∼90% specificity if biological replicate analysis is considered ground truth; see **Fig. S3c**). Together, these analyses show the high reproducibility of the strong genomic STARR-seq peaks.

### Genomic feature overlap analysis

Overlaps between the GP5d genomic STARR-seq and other genomic features were calculated from peaks called with MACS2^78^, with the exception of the TT-seq enhancers for which the estimated enhancer regions were used. Only chromosomes 1-22 & X were used in the analysis. All overlaps between the peaks were calculated using Bedtools^97^. The motifs used in calculating the matches to STARR-seq peaks are listed in **Table S5**. For comparison of STARR-seq peak overlap with ATAC-seq, H3K27ac ChIP-seq and TT-seq enhancers (**Fig. S7f,** upper part), 1 kb regions centered at peak summits instead of the peak boundaries were used in computing the Venn diagrams. Fisher’s exact test p-values were calculated using “bedtools fisher”^97^ after filtering out the positions in the ENCODE blacklist (peaks hitting these regions would not be considered).

Genomic STARR-seq in HepG2 cells was compared with data downloaded from the ENCODE project: ATAC-seq (ENCSR042AWH, replicate 1), histone modification ChIP-seq experiments for H3K27ac (ENCSR000AMO), H3K27me3 (ENCSR000AOL), and H3K9me3 (ENCSR000ATD), as well as ChIP-seq data sets for TP53 (ENCSR980EGJ), MED1 (ENCFF493UFO), and MED13 (ENCFF003HBS). The IDR-thresholded peaks and bigWig signal files showing fold change over control generated from all replicates were used except for ATAC-seq and TP53 ChIP-seq. The ATAC-seq replicate 1 reads were reanalyzed in hg19 coordinates in the same way as GP5d ATAC-seq data resulting in 60,500 peaks. For TP53 ChIP-seq, the ENCODE peaks were lifted over from GRCh38 to hg19 coordinates and replicate 1 reads were remapped to hg19 and MACS2 was used to generate a normalized coverage file. For the rest of the TFs and other chromatin-associated proteins the ChIP-seq peaks were taken from ref.^11^ (GEO accession GSE1042479), and the corresponding bigWig files were downloaded from the ENCODE portal. Super-enhancers for HepG2 are from http://www.licpathway.net/sedb.

The overlaps between different features were calculated using bedtools (chromosomes 1-22, X and Y). The Euler (**Fig. 2a**; R package eulerr) and bar (**Fig. S7g;** R package UpSetR) diagrams show other features overlapping the top quartile of all ATAC-seq and STARR-seq peaks according to maximum fragment coverage. When calculating overlaps between chromatin-associated proteins in open regions with or without STARR-seq signal (**Fig. S7e**), only STARR-seq peaks with IDR < 0.1 calculated from two STARR-seq replicates were used. The PolII-associated proteins POLR2AphosphoS2, PAF1, POLR2A, ZC3H4, TBP, POLR2AphosphoS5, and SSRP1 were excluded from this overlap analysis resulting in 202 proteins. In the overlap analysis, all Ensembl TSS (GRCh37, release 101) extended 1 kb to both directions were used.

### *De novo* motif mining

*De novo* motif mining for the peaks called from ATAC-seq, ChIP-seq and genomic STARR-seq were performed using HOMER^98^. Sequences used for motif mining of different enhancer classes in HepG2 cells were based on the intersections as in Euler/UpSet diagrams in **Fig. 2a; Fig. S7g**, with the overlaps calculated against top quartile of the ATAC or STARR-seq peaks, respectively. In addition, the sequences overlapping TSS-regions (from Ensembl GRCh37, release 101, extended 1 kb to both directions) were excluded, resulting in 1,524 closed chromatin enhancers, 971 classical enhancers, and 3,797 chromatin-dependent enhancers that were used in the analysis. For the sequences enriched from the random enhancer and random promoter-random enhancer STARR-seq experiments and from the ATI assay, *de novo* motif mining was done using the “Autoseed” program as described earlier^29^. From the random enhancer STARR-seq in GP5d cells, the sequences were cut to 40 bp long non-overlapping subsequences starting from position 6 and ending at position 165, and the subsequences containing N were removed (see **Table S7** for the numbers of analyzed subsequences). “Autoseed” program was also used to mine *de novo* motifs enriched at specific positions in relation to TSS using TSS-aligned sequences from GP5d cells and input DNA sequences sampled from the same positions (**Table S7**). The resulting seven *de novo* motifs were mined from the subsequences spanning the following positions in relation to TSS: two initiator motif variants from position one at TSS (only forward strand), CREB motif and two CREBMAF heterodimer variants from -30 to 30, and TATA box promoter from -30 to 23. These motifs were also included to motif enrichment comparison between promoter and enhancer sequences.

### Conservation of genomic STARR-seq elements

Conservation of genomic STARR-seq peaks and input fragments was analyzed by calculating their average GERP scores^99^ using precomputed base-wise GERP scores (hg19) from ref.^100^ (http://mendel.stanford.edu/SidowLab/downloads/gerp/). In the analysis, only chromosomes 1-22 were included, and the elements overlapping ENCODE blacklisted regions (ENCFF419RSJ) and UCSC RepeatMasker^101^ “Repeats” track were discarded. A base pair was deemed conserved if its GERP score was higher than the average GERP score (∼2.2) of the coding sequence^100^. Additionally, the GERP scores were calculated for three known enhancers as detailed below.

- GP5d genomic STARR-seq peaks, mean GERP score = 0.14. The average number of conserved base pairs in 170 bp surrounding the STARR-seq peak summits was ∼50.8 (∼169.4 for whole peaks, corresponding to ∼42.6 when the average width of a STARR-seq peak, ∼675.5 bp, is scaled to 170 bp).
- 100,000 randomly sampled genomic STARR-seq input fragments, mean GERP score = 0.01.
- The MYC335 enhancer (chr8: 128413174-128414429)^102^, mean GERP score ≈ 3.07, number of conserved base pairs = 921.
- The SHH enhancer (chr7: 156583796-156584568)^103^, mean GERP score ≈ 3.66, number of conserved base pairs = 642.
- The Sox9 enhancer (chr17: 69480826-69481362)^104^ with respective coordinates for human SOX9 enhancer (hg19) obtained using UCSC genome LiftOver tool (https://genome.ucsc.edu/cgi-bin/hgLiftOver), mean GERP score ≈ 0.70, number of conserved base pairs = 230.

### Preprocessing of the random promoter-enhancer pairs

The STARR-seq enhancer sequences derived from RNA were mapped to corresponding promoter-enhancer pairs in the input DNA by exact matches of the first 20 bases of the 150 bases long enhancer sequences. Duplicate sequences were removed as described for random enhancers, except that three 40 bases long subsequences from 16 to 135 were used thus ensuring that only one of the sequences with Hamming distance less than 3 was chosen. Then, promoter and enhancer sequences were filtered separately by removing 1) all adapter sequences that included some (partial) adapter sequence according to cutadapt^105^, 2) sequences that mapped to plasmid backbone sequence using bowtie2^77^, and 3) outlier sequences in terms of nucleotide composition (count of any nucleotide more than 3 median absolute deviations higher than the median count). Input DNA sequences were processed the same way. For promoter-enhancer pair analyses, the remaining promoter-enhancer pairs were collected and pairs containing highly similar sequence as promoter and enhancer were removed. The numbers of sequences used in downstream analysis are shown in **Table S7**.

### Mapping TSS positions based on template switching

First, sequences derived from spliced transcripts were identified using the constant sequence spanning the splice site after intron removal (cutadapt program); other sequences were not processed further. Next, UMI sequence was removed from the 5’ end of each sequence and the last 20 bp of its random part was used to recognize the corresponding promoter from the input DNA. To accurately recognize the first base of the transcript and thus the position of the TSS, it was assumed that the template switch process had added at least 3 and at most 4 guanines to the 5’ end of the transcript. On this basis, only the RNA sequences starting with at least three Gs were used in the analysis. Each such sequence was aligned to the corresponding input DNA promoter sequence using an exact 20 bp match starting from the 6th base to allow for the extra Gs. Finally, the Gs added by the template switch were trimmed and discordant sequences removed according to the alignment. The frequency of the four Gs instead of three was estimated from the sequences that do not have G at the fourth position in the alignment to the input. For those that did have a G also in the input sequence, removing three or four Gs was decided randomly but so that the frequency of the fourth G matched the estimate. The two GP5d template switch libraries were processed separately and then merged so that only one transcript was kept for each input DNA promoter sequence to prevent duplicate promoter sequences. The exact positions of the TSS at the promoters were recorded and the flanking sequences were used for further analysis. The numbers of sequences obtained are listed in **Table S7** as the number of flanking sequences fitting to the random region depends on the flank sizes. The comparison to human endogenous promoters was done using TSS positions from EPD database (hsEPDnew 006).

### Matching of known motifs

The motifs were matched to sequences using MOODS^106^. The matching was done separately for each strand using p-value thresholds (1 × 10^-6^, unless otherwise stated) and strand- and sample-specific nucleotide frequencies. When calculating motif matches for logistic regression PWM features, affinity threshold of 0 was used instead of p-value. Motif matches that resulted in occupancy probabilities smaller than 0.01 were discarded (see Logistic regression classification for details). To determine the activity of an individual motif, the total number of its matches in the sequences was counted so that overlapping matches in different strands were counted only once. Then fold changes between the motif match counts in RNA and a randomly sampled subset of input DNA were estimated using the function PsiLFC in R package lfc. When comparing two different RNA samples (for example two replicates), two different random samples of input DNA were used to avoid overestimating similarities by using identical input counts. To estimate the effect of the number of binding sites in the same sequence, the number of sequences having exactly two non-overlapping motif matches was counted and the fold change was compared to the fold change of those having exactly one match. If the site occurrences are independent of each other, the expected frequency of several sites is the product of the individual frequencies. Thus, the expected log_2_ fold change assuming independent actions of several motif occurrences was calculated as the sum of their individual log_2_ fold changes.

### Analysis of interactions between promoter and enhancer

Both for RNA and randomly sampled input DNA promoter-enhancer pairs, the number of such pairs that one motif occurs in the promoter and a second one in the enhancer was counted for each motif pair (excluding heterodimers) and motif match strand combination (++, +-, -+, --). The counts over the strand combinations were summed to get the total number of pairs and the fold change of the number the pairs between input DNA and RNA was estimated using the function PsiLFC in R package lfc. If the promoter and enhancer occurrences are independent of each other, the expected frequency of the pair of sites is the product of the individual frequencies. The expected log_2_ fold change assuming independent actions of the promoter and enhancer motif was thus calculated as the sum of their individual log_2_ fold changes. The same analysis was done using the reversed but not complemented control motifs.

### Motif match positioning relative to TSS and STARR-seq vector

Motifs were matched to TSS flanking promoter sequences and for each motif only the highest affinity match per sequence was considered. The number of matches for each motif was then counted separately at each position and strand. To get positional activity scores for position-specific regression analysis, motif matching was done for TSS flanking sequences from position -100 to +20 in relation to TSS and for a control set generated by sampling for each TSS sequence a subsequence of same length from the same position from an input DNA promoter (p-value threshold 5 × 10^-4^). The log_2_ fold changes of the motif match counts between TSS flanking set and control set (estimated with the lfc package) were then used as a positional activity score for each position and strand.

To study p53 motif match positioning relative to the STARR-seq vector, the motif was matched (p-value threshold 10^-5^) to highly selected sequences chosen by taking only sequences observed at least twice in both GP5d enhancer replicate experiments. A histogram of match start positions was generated by counting only the highest affinity match in each sequence. A smoothed density estimate was generated using a gaussian kernel (R ggplot geom_density with adjust=0.5).

### Mutual information (MI) analysis

The binding events on the aligned STARR-seq reads (40+40 bp surrounding TSS) were analyzed by calculating the mutual information (MI) between 3-mer distributions at two non-overlapping positions of the aligned sequences. MI can be used to capture binding events, because if the binding contacts two continuous or spaced positions (3-bp wide) on the sequences at the same time, correlations will be observed for the 3-mer distributions at the two positions. Subsequently, the biased joint distribution will be detected as an increased MI between the positions. For two non-overlapping positions (pos1, pos2), the MI between them was estimated as reported previously^41^. The calculation uses the observed frequencies of a 3-mer pair (3+3- mer), and of its constituent 3-mers at both positions:

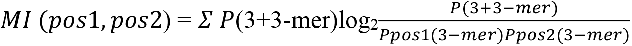

where *P(3+3-mer)* is the observed probability of the 3-mer pair (i.e. gapped or ungapped 6 mer). *P_pos1_(3-mer)* and *P_pos2_(3-mer)*, respectively, are the marginal probabilities of the constitutive 3-mers at position 1 and 2. The sum is calculated over all possible 3-mer pairs using 50,787 TSS sequences. When calculating the positional 3-mer and the pairwise 3+3-mer distributions, a pseudocount of 10 was added to each k-mer.

### Data preprocessing for machine learning analysis

The datasets used in each machine learning analysis and their division into training, test and validation sets are detailed in **Table S7**. To enable sequences from genomic measurements (genomic STARR-seq and ATAC-seq) to be scored on the CNNs that were trained on the random enhancer STARR-seq data and vice versa, the length of the sequences fed to these models was standardized to 170 bp. Thus, additional preprocessing specific to machine learning analyses was done for the genomic STARR-seq and ATAC-seq data.

First, an extended blacklist file was created to remove possibly problematic genomic regions that might cause the machine learning models to learn biases instead of real signals. In addition to the standard ENCODE blacklist (ENCFF419RSJ), this extended blacklist contains all positions ±1Mb from centromeres, all positions with Ns in the hg19 reference genome and non-uniquely mapping regions that were defined as follows: All unique 55-mers present in the hg19 reference genome were fetched and aligned back to hg19 reference with bwa aln^84^ algorithm. Then each position that was not covered by reads mapped with MAPQ>20, was added to the extended blacklist. This extended blacklist covers around 12% of the hg19 reference genome.

GP5d genomic enhancer fragments were created by fetching the sequence (hg19) that maps between the paired end reads. The signal set (class 1) sequences were created by taking the 170 bp closest to the peak summit from each genomic STARR-seq fragment that overlaps with a GP5d genomic enhancer STARR-seq peak. Balanced control sets of 170 bp sequences (class 0) were sampled from the genomic STARR-seq input requiring that the reads map and do not overlap with regions covered by the extended blacklist or with GP5d genomic enhancer STARR-seq peaks. To ensure that the classifier does not learn any features possibly correlating with different input library coverage between the class 1 and class 0 sequences, the class 0 sequences were sampled in such a way that their input library coverage histogram matched the input library coverage histogram of the class 1 sequences. Input library coverage of each class 1 and class 0 sequence was calculated by counting how many input library fragments each of them overlaps. Then 36 evenly sized bins were created so that the first bin included coverages between 0 and 9 and the last between 350 and 359 (354 was maximum coverage for class 1 sequences). Then class 0 sequences were sampled so that their count in each bin equaled the count of class 1 sequences in that bin for each set (training, test, validation) separately. Sequences from random STARR-seq experiments were not mapped to any reference sequence at any point, so this precaution is not relevant with random STARR-seq experiments.

The GP5d ATAC-seq single-end short reads were extended to the average fragment size of the library (300 bp) by adding 300-readlen to 3’-end of each read (where readlen is the length of each read). For the class 1 signal set, extended fragments that overlap with any ATAC-seq peak were selected and the 170 bp sequence closest to the overlapping peak summit was retrieved from the fragment. Exact duplicate sequences were discarded. A balanced negative set (class 0) was created by sampling random 170 bp sequences from the genome and not allowing them to overlap with ATAC-seq peaks or regions covered by the extended blacklist.

### Logistic regression classification

The logistic regression classifiers were implemented using the LogisticRegression function from scikit-learn library^107^. All logistic regression models were regularized with L1 norm. Using L1 norm as regularization is important for the interpretability of the model coefficients, as the set of PWMs used as features of the model (**Table S10**) contains several matrices that can be very similar to each other. Thus, a non-regularized model could split effects into coefficients of similar PWMs. Using the L1 norm that penalizes solutions with higher number of non-zero coefficients enables finding the best performing model with the lowest number of individual PWMs contributing to the model. The optimal regularization strength was chosen based on the area under precision-recall curve on the validation data (see **Tables S8, S9** for the tested and final values, respectively). Classification performance on unseen test data is shown for each trained hyperparameter combination in **Fig. S9h**. Otherwise the regression was run on default parameters. First, a logistic regression classifier was trained using only features that count matches of individual PWMs (880 features, **Table S10**). After this, a more complex classifier was fit with additional features counting all self-pairs (*A_i_+A_i_*, *i* runs over all the 880 PWMs), and all pairs of the top 20 strongest individual features (20 features with largest absolute value of the regression coefficient) from the simpler model (**Table S11**) with all other PWMs (*S_j_+A_i_*, *i* runs over all the 880 PWMs, *j* runs over the top 20 PWMs from the simple model of 880 features). Thus, this more complex model is expected to cover all meaningful general pairwise interactions between the binding motifs of HT-SELEX PWMs.

For each feature (corresponding to a PWM, *X*), the probability that a given read is occupied by a given TF or TF-pair is calculated by using an approach derived from ref.^108^, where:

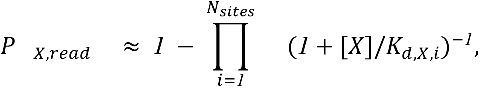

where *N_sites_* is the number of motif matches for PWM *X* in the read, *[X]* is the free concentration of the TF corresponding to the PWM *X* and *K*_*d,x,i*_ = *exp*(−*ΔG*_*x,i*_/*RT*) is the equilibrium dissociation constant of the binding site *i* of the TF corresponding to the PWM *X*. In ref.^108^, the free concentration of each TF was set to equal the *K_d_* of the consensus sequence. However, for some TFs with a long PWM, exact matches to the consensus sequence are rare, and setting the scoring as described above will result in the occupancy scores 0 for many functional binding sites, reducing the variance of the scores of the variables corresponding to these TFs. To overcome this, we used a normalization approach based on the fact that TFs generally have ∼10000-300000 binding sites in the human genome and defined the free concentration of each protein to correspond to the *K_d_* of the strength of its 10000th strongest binding site in the human genome. Thus,

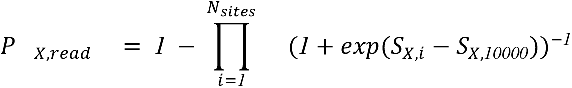

where PWM match score of the 10000th strongest match of the PWM *X* is *S_X,10000_* and the PWM match score of the ith site in a read is *S_X,i_*, This normalization was calculated separately for each PWM and strand. The probability for a pair of TFs (X+Y) to occupy a sequence^108^ is

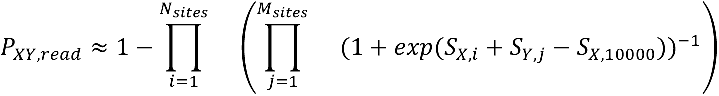

### Position specific logistic regression classification

The positional activity scores of the 880 dominating set PWMs and the 7 core promoter PWMs (see **Table S12**) were used to train a position-specific logistic regression classifier on the promoter capture STARR-seq data. In contrast to the simple logistic regression described above, the PWM match scores were weighted using the positional activity scores of the corresponding PWM. Thus, instead of occupancy probabilities, scores

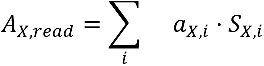

were calculated for each sequence and PWM feature *X*, where *a_X,i_* is the positional activity score of PWM *X* at position *i* and *i* runs over all matches of PWM *X* in a sequence. Regression coefficients were learned separately for both strands for each PWM. Training of the position-specific regression model was done similarly to the training of other logistic regression models. Final model was selected based on area under precision-recall curve on separate validation data. Classification performance on unseen test data is shown for each trained hyperparameter combination in **Fig. S9h**.

### Convolutional neural network classification

The CNN classifiers used the raw fasta sequences as input to learn the features during the training process. The random enhancer CNN was trained using also the reverse complement sequences of the training data, all other models were trained using one orientation only. The CNN models consist of convolutional modules with a 1D convolutional layer followed by batch normalization, ReLu activation and a dropout layer. The number of convolutional modules, the number of convolutional filters per layer, and the dropout rate are hyperparameters that were optimized based on the validation data. The convolutional modules used dilated convolution^109^ so that the dilation rate of the *i*th layer is *i*^2^, which allows learning interactions with fewer parameters than fully connected (dense) layers. After the convolutional modules, the final layer is a dense layer of two nodes with sigmoid activation. All the machine learning models used in this work will be available at Zenodo (https://zenodo.org/). This CNN architecture was selected by testing it against architectures with varying sized dense layers after the convolutional layers. Adding the dense layers did not improve the performance of the models. The models were built on Keras (https://keras.io/) using TensorFlow 1.14.0 backend^110^.

Models were trained using the Adam optimizer with default parameter values. Training was stopped once binary accuracy on validation data did not improve within 200 epochs or when the total training time on a single Nvidia Volta V100 GPU exceeded 72 hours (an exception being the “double input” CNN models trained to classify the “binary STARR-seq” data (see below for more details) where training was continued up until 144 hours if needed). Model parameters were initialized using the He uniform variance scaling initializer^111^. All models, except the double input CNN, were trained on batch size 128 as changing the batch size had a negligible effect on classification performance, and only had one convolutional body and no dense layers (except the final output layer). Batch size for the double input CNN models was determined over grid search between 32 and 64. The other hyperparameters of the CNN models were determined by grid search over the values shown in **Table S8**, and the hyperparameters of the final selected models are shown in **Table S9**. Classification performance on unseen test data is shown for each trained hyperparameter combination for each trained CNN model in **Fig. S9h.**

### Note on optimal classification of random enhancer STARR-seq data

As seen from **Fig. S9h**, classifying between random enhancer STARR-seq signal and input sequences is more difficult than for other datasets studied here. In addition to the logistic regression and CNN classifiers described in detail here, also random forest classifiers and support vector machines trained on PWM match data similarly to the logistic regression model failed to classify random enhancer STARR-seq data better than the CNN model (data not shown). We believe that this limited classification accuracy is mainly a feature of the experimental design. Transcription of one molecule of DNA is an inherently probabilistic single-molecule process, and therefore some transcription occurs at random. Some sequences are thus recovered due to this “transcriptional noise”, limiting the maximum classification accuracy. To determine whether a better classification is possible, the replicate experiment of the GP5d random enhancer STARR-seq was utilized to create an unseen test set that contains only those original test set sequences that were also observed in the replicate experiment (2.8% of sequences were observed in both replicates). A balanced number of input library sequences were used as class 0 sequences. Using the final GP5d random enhancer STARR-seq CNN model on this high-confidence test set resulted in ∼4% increase in AUprc (**Fig. S4d**). The mean(AUprc) ± std from eight CNN models with different hyperparameters was AUprc_all_=0.614±0.004 for test set with all sequences and AUprc_common_=0.641±0.003 for the common sequences between the replicates.

We also further removed the sequences observed more than once in the input library from the 371,390 common STARR-seq sequences between the GP5d random enhancer STARR-seq replicates to evaluate whether possible uneven sequence representation in the input library could affect the classification. For each common STARR-seq library sequence between replicates, the number of matches in the input library was counted by requiring the first 60 bases of the common sequence to match exactly to the first 60 bases of a read 1 sequence in the input library. There were only 6,213 (∼0.02%) common sequences between replicates that were observed more than once in the input library (total 999,854,466 input sequences). Removing these 6,213 sequences had a negligible effect on the classification (**Fig. S4d**).

As an external evaluation we trained the previously published gapped k-mer SVM classifier^112, 113^ on the high-confidence GP5d random enhancer STARR-seq data where the signal set (class 1) sequences were observed in both of the replicates. A balanced set of class 0 sequences were sampled randomly from the input library. We used the default k-mer length (11) and the default decay strength for the center weighted gkm kernel, and tested both “gapped k-mer”, and “gapped k-mer + center weighted” kernels. We optimized the regularization strength over values (1, 0.1, 0.01, 0.001) with the validation data (by calculating area under precision-recall curve). The best hyperparameter combination was obtained by using gapped k-mer kernel and regularization strength=0.1, producing AUprc=0.6207 on the validation data. With the unseen test set (high-confidence sequences) this model obtained AUprc=0.6086. With the full test set from GP5d random enhancer STARR-seq replicate 1 the gapped k-mer SVM obtained AUprc=0.57, which is slightly better than the pairwise logistic regression (0.56) but worse than the CNN (0.62). The gapped k-mer SVM training with the full GP5d random enhancer STARR-seq training set of 13,237,548 sequences did not complete within approximately three weeks of running time on our computational cluster with 16 parallel threads (gapped k-mer SVM maximum), so performance of a model trained with the full data set could not be evaluated.

### Differential expression prediction

A lasso regression model was used to study the extent to which differential gene expression between GP5d and HepG2 cell lines could be predicted using STARR-seq and ATAC-seq derived features. Logarithmic fold change between GP5d and HepG2 expression values (tpm) was used as the target variable for regression (see section “Chromatin immunoprecipitation (ChIP-seq) and gene expression analysis (RNA-seq and CAGE)” above). The STARR-seq and ATAC-seq peaks were divided into 12 explanatory features based on the cell line in which the peaks were observed and the information about whether the ATAC-seq peaks were promoter proximal (less than 1kB distance to any gene in the target gene set) or not as follows: Common.ATAC.enh.noSTARR; Common.ATAC.enh.yesSTARR; Common.ATAC.prom; Common.STARR.noATAC; GP5d.ATAC.enh.noSTARR; GP5d.ATAC.enh.yesSTARR; GP5d.ATAC.prom; GP5d.STARR.noATAC; HepG2.ATAC.enh.noSTARR; HepG2.ATAC.enh.yesSTARR; HepG2.ATAC.prom; HepG2.STARR.noATAC. Naming “Common.STARR.noATAC” means STARR-seq peaks that were observed in both cell lines and that did not overlap with an ATAC-seq peak, and “HepG2.ATAC.enh.yesSTARR” means HepG2-specific ATAC-seq peaks that are not within 1 kb from target gene TSSs and do overlap with STARR-seq peaks.

For each feature, the logarithmic fold change at peak summit reported by MACS2 (LFC) was used as the strength of the peak. Peak summit position was used as the position of the feature (ATAC-seq peak summits were used for all other features except the ones that had no overlap with ATAC-seq). Peak score S, meaning the effect of a peak to a TSS that is d bp away from the peak summit for each feature was calculated assuming it decays like:

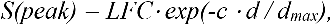

where LFC is the logarithmic fold change at peak summit, c is a scaling parameter and d_max_ is the maximum distance of a peak from the TSS for its effect to be included in the model. This peak score S was used to quantify the effect of each feature in the regression model.

The target genes were split into training (8815 genes), validation (1449 genes) and test (2321 genes) sets according to chromosomes they are in (training: chr1, chr3, chr5, chr7, chr9, chr11, chr13, chr14, chr15, chr16, chr17, chr18, chr19, chr20, chr21, chr22, chrX; validation: chr4, chr6, chr8; test: chr2, chr10, chr11). A lasso regression model was trained using the LassoCV method in scikit-learn^107^ library (version 0.24.1) where the regularization strength (L1 norm) was determined using 5-fold cross-validation during training (n_alphas=100, no intercept term). On top of this, the model hyperparameters c and d_max_ were optimized using the validation data set with a grid search over the values shown in **Table S8**. Features were standardized by removing the mean and scaling to unit variance before model training. The best model had the following hyperparameters: d_max_=100,000 bp; c=4.3; regularization strength α=0.013203848400344683. This means that the effect of a peak decays to ½ at approximately 16,120 bp from the peak summit. This model obtained R^2^ = 0.1227036350306634. Regression coefficients of this best performing model are shown in **Table S6**.

### Interpretation of convolutional neural network classifiers

To test which HT-SELEX PWMs the convolutional neural network had learned, the trained CNN was used as an “oracle” as in ref.^66^ by embedding 100 sequences drawn randomly from each PWM to random enhancer STARR-seq input library sequences (**Table S10**). Each such sequence was embedded at a random position to one of 100 different randomly chosen input sequences (same input sequences used as background for each PWM) and the average enhancer probability over the 100 sequences was calculated for each PWM. When embedding a single PWM per input sequence, first, the position for the embedding was drawn from random uniform distribution. Next, the embedded sequence was drawn by random from the corresponding PWM. When embedding a motif pair, first the positions of the embedded sequences were drawn at random not allowing overlap between the embedded sequences. Then, both of the embedded sequences were drawn independently from the corresponding PWMs. Thus, both the positions and the distance between the embedded sequences are random. The expected enhancer probability for a sequence with two embedded PWMs given that there are no interactions between them is *p*_2_ = *1* − (*1* − *p_1_*)^2^, where *p_i_* is the enhancer probability of a sequence with *i* PWMs embedded, representing the cumulative probability for geometric distribution with two trials.

The TERT promoter sequences were visualized with the DeepLIFT software^73^, which shows the importance of each position in a sequence for the final prediction verdict. Positive values correspond to sites that move the prediction verdict towards “promoter” class and negative values to sites that move the verdict towards “not promoter” class. The activation values from the wild type and mutated promoter sequences were compared against 15 randomly selected background sequences from the random promoter STARR-seq input and their average signals were visualized. Predicted promoter probabilities of the TERT promoters were obtained by scoring the sequences with the CNN trained on promoter capture STARR-seq data.

To visualize pairwise position-specific interactions learned by the CNN classifiers trained on STARR-seq and EPD promoters, respectively, the high-confidence promoter sequences were visualized with the same mutual information (MI) analysis pipeline^114^ as described above. Here, we generated 10 million sequences of length 120bp from random uniform nucleotide background and scored each of them with the 10 best (according to binary accuracy on the validation data) STARR-seq promoter capture CNN models and 10 best EPD promoter models. Those sequences obtaining a promoter probability of 0.9 or higher according to each of the 10 models were selected for MI analysis. This resulted in 51,131 sequences for STARR-seq promoter capture CNN and into 395,663 sequences for EPD promoter CNN. These numbers together with the MI-plot themselves in **Fig. S9f,g** indicate that the STARR-seq data allowed learning models with stricter positional dependencies.

### Validation of the predicted promoter mutation effects with external data

To validate the effect of mutations predicted by the CNN model trained on promoter capture STARR-seq data, the model predictions were correlated with a saturation mutagenesis study of the same promoter^46^ (see **Fig. S9b-d**). The statistical significance of the mutation effects predicted by the CNN model was estimated as follows. First, the predicted promoter probabilities from the CNN model were transformed into log odds scores logit(p)=log(p/(1-p)), and the predicted effect of each mutation was calculated as the logarithm of odds ratio between the predicted promoter probability of the mutated sequence and the wild type sequence, namely

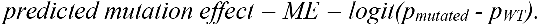

Then, an empirical p-value was calculated for each predicted TERT promoter mutation effect by comparing if the predicted effect in TERT promoter is bigger than predicted effect on shuffled promoter sequences at the same position and for the same type of mutation. This was done by creating 10,000 shuffled sequences (dinucleotide frequencies preserved) of the TERT wild type promoter. For each of these, all possible SNPs were introduced, and the predicted mutation effects were calculated for each of them as the logarithm of odds ratio against the predicted promoter probability of the wild type (shuffled) promoter sequence. For each position and mutation type, the empirical p-value was calculated as the fraction of the predicted mutation effects on the shuffled sequences that were more extreme than the predicted effect on the wild type TERT promoter:

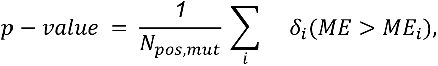

for cases where the predicted promoter probability increases due to the mutation and

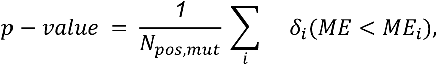

for cases where the predicted promoter probability decreases due to the mutation. *N_pos,mut_* is the total number of shuffled sequences where mutation *mut* was observed at position *pos* and the summation index *i* runs over all such sequences, δ_*i*_(*ME<ME_i_*) is 1 if *ME<M_i_* and otherwise 0. All mutations that were non-significant according to either the empirical p-value or the p-value from the saturation mutagenesis experiment from ref.^46^ were filtered out (threshold p-value<0.05). The correlation of the model predictions between saturation mutagenesis logarithmic fold changes is shown in **Fig. S9b-d**. The saturation mutagenesis experiments used here are TERT-HEK (HEK293T cells) and TERT-GBM (primary glioblastoma SF7996 cells).

### Promoter-enhancer interaction analysis using machine learning

The “binary STARR-seq” design allows looking for relatively short-range interactions between promoters and enhancers. To comprehensively test for any such interactions, we trained several models with a double input CNN architecture, where the promoter and the enhancer parts of the sequence are read in with separate input convolutional bodies (a convolutional body consists of multiple layers of convolutional modules with dilated convolutions) and later connected with a fully connected layer to learn interactions between the outputs of the two convolutional bodies. Depth of the convolutional bodies was optimized with separate validation data along with other hyperparameters detailed in **Table S8**. The rationale of this design is that the convolutional modules with dilated convolutions are capable of learning interactions within the promoters and within the enhancers, while the subsequent fully connected layer then integrates information between the promoters and the enhancers.

The search for interactions between promoters and enhancers was implemented by shuffling the training data of the models and keeping the model architecture constant, We trained double input CNN models with four different types of training data: i) The paired readout from the binary STARR-seq experiment with all possible information about interactions between promoters and enhancers intact, referred to as “binary STARR-seq CNN (paired)” in **Fig. 5c**. ii) Permutated training data, where the pairing between the promoter and the enhancer sequences is broken so that any specific interactions between the promoters and enhancers are killed, but all promoters and enhancers come from promoter+enhancer pairs that are active, referred to as “binary STARR-seq CNN (permutated)” in **Fig. 5c**. iii) Training data where only the promoter comes from active promoter+enhancer pairs and the enhancer sequences are sampled randomly from the inactive input pool of enhancer sequences, referred to as “binary STARR-seq CNN (enhancer from input)” in **Fig. 5c**. iv) Training data where only the enhancer comes from active promoter+enhancer pairs and the promoter sequences are sampled randomly from the inactive input pool of promoter sequences, referred to as “binary STARR-seq CNN (promoter from input)” in **Fig. 5c**.

### Transcription start site prediction

All the promoter models were trained on data where the TSS is 100 bp from the start of the training sequence. Thus, scoring any 120 bp sequence with these models gives a probability that the position 100 in these sequences is a TSS of a functional promoter sequence. Each possible TSS position within ±500 bp from the TSSs of the active GP5d promoters (and separately of all EPD promoters) was analyzed by taking 100 bp upstream and 20 bp downstream from the candidate TSS and by scoring these sequences with the promoter models. For each test set active GP5d TSS and for each promoter model, the position obtaining the highest promoter probability from the corresponding model was taken as the predicted TSS position.

### Preprocessing of genomic promoters

Human promoter coordinates were obtained from the eukaryotic promoter database^47^ (EDP, version 006, hg19) and sequences 100 bp upstream and 20 bp downstream of the TSS were fetched. Promoters overlapping with the extended blacklist described earlier or residing outside of chromosomes 1-22 or X were discarded. The division to training, test and validation sets for machine learning is detailed in **Table S7**. The genomic promoter control set (class 0) was generated by randomly drawing a balanced number of 120 bp sequences (according to the same training, test and validation split) that do not overlap with EPD promoters or regions in the extended blacklist described above. For cancer-associated mutation analysis, the genomic sequence 100 bp upstream and 20 bp downstream of the TSS for TERT transcript ENST00000310581.5 was downloaded from Ensembl GRCh37 release 99 Biomart. The top three mutation hotspots in the sequence as well as the recurring mutations (mutations observed in more than one patient) were obtained from ref.^45^.

### CAGE analysis

The 5’ ends of the GP5d CAGE reads contain a 3-bp barcode (ATC) followed by a 6 bp constant sequence (CAGCAG). In addition to removing these, the next 2 bp that were mostly Gs, according to a FastQC (https://www.bioinformatics.babraham.ac.uk/projects/fastqc/) quality control report, were discarded. The reads were aligned to a combined Phi × 174 + hg19 reference genomes using bwa mem^115^. Only the reads mapping to hg19 with MAPQ 30 or higher were extracted and duplicates were removed with samtools rmdup. The active promoters were discovered from the mapped CAGE reads by clustering the 5’-ends of the reads with paraclu software^116^. In total, paraclu called 7365 clusters (peaks) from the GP5d CAGE data fulfilling the following criteria: 1) Cluster is supported by more than 9 unique reads. 2) Cluster cannot be longer than 200 bp. 3) Remove clusters where the maximum 5’ end density per base divided by the baseline density is less than 2. 4) Remove any cluster that is contained within a larger cluster. The active GP5d promoters, used as the test set in predicting TSS position, were defined as those EPD test set (see **Table S7**) promoters that overlap with a GP5d CAGE peak. Thus, the TSS positions come from EPD.

## Data and code availability

All sequence data is available under ENA accession xxxxx, and custom code will be made available upon request. All pre-trained machine learning models are available at Zenodo with accession xxx. Training, test and validation data sets for the CNN models are available at Zenodo with accession yyy.

## Acknowledgements

We thank Dr. Minna Taipale for critical review of the manuscript, and Anu M. Luoto, Kaisu Jussila, Åsa Kolterud for technical assistance. We also thank HiLIFE research infrastructures including Biomedium Functional Genomics (FuGU), FIMM technology center and sequencing core facilities of Karolinska Institute and SciLife lab. We wish to acknowledge CSC – IT Center for Science, Finland for computational resources. We also thank the Eukaryotic promoter database for the publicly available datasets (https://epd.epfl.ch//index.php). This work was supported by grants from Academy of Finland (Finnish Center of Excellence program: 2012-2017, 250345 and 2018-2025, 312041, Post-doctoral fellowships; 274555, 288836 and Research Fellowships, 3177807), Finnish Cancer Foundation, research grants to BS from Jane and Aatos Erkko Foundation and personal grant to TH from Emil Aaltonen Foundation. PC was supported by the Deutsche Forschungsgemeinschaft within SFB860 and SPP1935 and under Germany’s Excellence Strategy (EXC 2067/1-390729940), the European Research Council Advanced Investigator Grant TRANSREGULON (grant agreement No 693023), CIMED and SciLifeLab.

## Author contributions

JT conceived and supervised the study. BS designed all the custom CpG-free STARR-seq vectors, cloning and custom Illumina sequencing strategy for generation of libraries from TF motifs, genomic DNA and random sequences and performed all the STARR-seq experiments with help from PP. BS performed the ATAC-seq, ChIP-seq, RNA-seq, template switch PCR libraries, and CRISPR-Cas9 edited TP53-null GP5d cell lines. BS performed the processing of Illumina sequencing data, peak calling and de novo motif analysis for STARR-seq, ATAC-seq, ChIP-seq data. TK designed the synthetic TF motif libraries, performed all STARR-seq analysis from all different designs from motif, genomic DNA, random DNA and template switch libraries. TH performed the logistic regression, CNN classifiers, training models, machine learning analyses, mutual information analyses, conservation and overlap analysis of genomic enhancers and processing of CAGE and promoter datasets. BW performed ATI and CAGE data were from KD and CD. FZ helped with the MI plot and EK helped with the ATAC-seq pipeline. KL, ML and PC contributed the TT-seq data. BS, TH, TK, PP and JT wrote the manuscript with input from all authors.

## Competing interests

The authors declare no competing interests.

**Supplementary information** is available for this paper.

